# The combination of Hebbian and predictive plasticity learns invariant object representations in deep sensory networks

**DOI:** 10.1101/2022.03.17.484712

**Authors:** Manu Srinath Halvagal, Friedemann Zenke

## Abstract

Discriminating distinct objects and concepts from sensory stimuli is essential for survival. Our brains accomplish this feat by forming disentangled internal representations in deep sensory networks shaped through experience-dependent synaptic plasticity. To elucidate the principles that underlie sensory representation learning, we derive a local plasticity model that shapes latent representations to predict future activity. This Latent Predictive Learning (LPL) rule conceptually extends Bienenstock-Cooper-Munro (BCM) theory by unifying Hebbian plasticity with predictive learning. We show that deep neural networks equipped with LPL develop disentangled object representations without supervision. The same rule accurately captures neuronal selectivity changes observed in the primate inferotemporal cortex in response to altered visual experience. Finally, our model generalizes to spiking neural networks and naturally accounts for several experimentally observed properties of synaptic plasticity, including metaplasticity and spike-timing-dependent plasticity (STDP). We thus provide a plausible normative theory of representation learning in the brain while making concrete testable predictions.

---

Recognizing invariant objects and concepts from diverse sensory inputs is crucial for perception. Watching a dog run evokes a series of distinct retinal activity patterns that differ substantially depending on the animal’s posture, lighting conditions, or visual context (Fig. 1a). If we looked at a cat instead, the resulting activity patterns would be different still. That we can effortlessly distinguish dogs from cats is remarkable. It requires mapping *entangled* input patterns, which lie on manifolds that “hug” each other like crumpled-up sheets of paper, to *disentangled* neuronal activity patterns, which encode the underlying factors so downstream neurons can easily read them out [1]. Such transformations require deep sensory networks with specific network connectivity shaped through experience-dependent plasticity (Fig. 1b). However, current data-driven plasticity models fail to establish the necessary connectivity in simulated deep sensory networks. At the same time, supervised machine learning algorithms *do* yield suitable connectivity [2] in deep neural networks (DNNs) that further reproduce essential aspects of the representational geometry of biological neural responses [3, 4]. This resemblance proffers DNNs as potential tools to elucidate neural information processing in the brain [5, 6].

**Figure 1:**
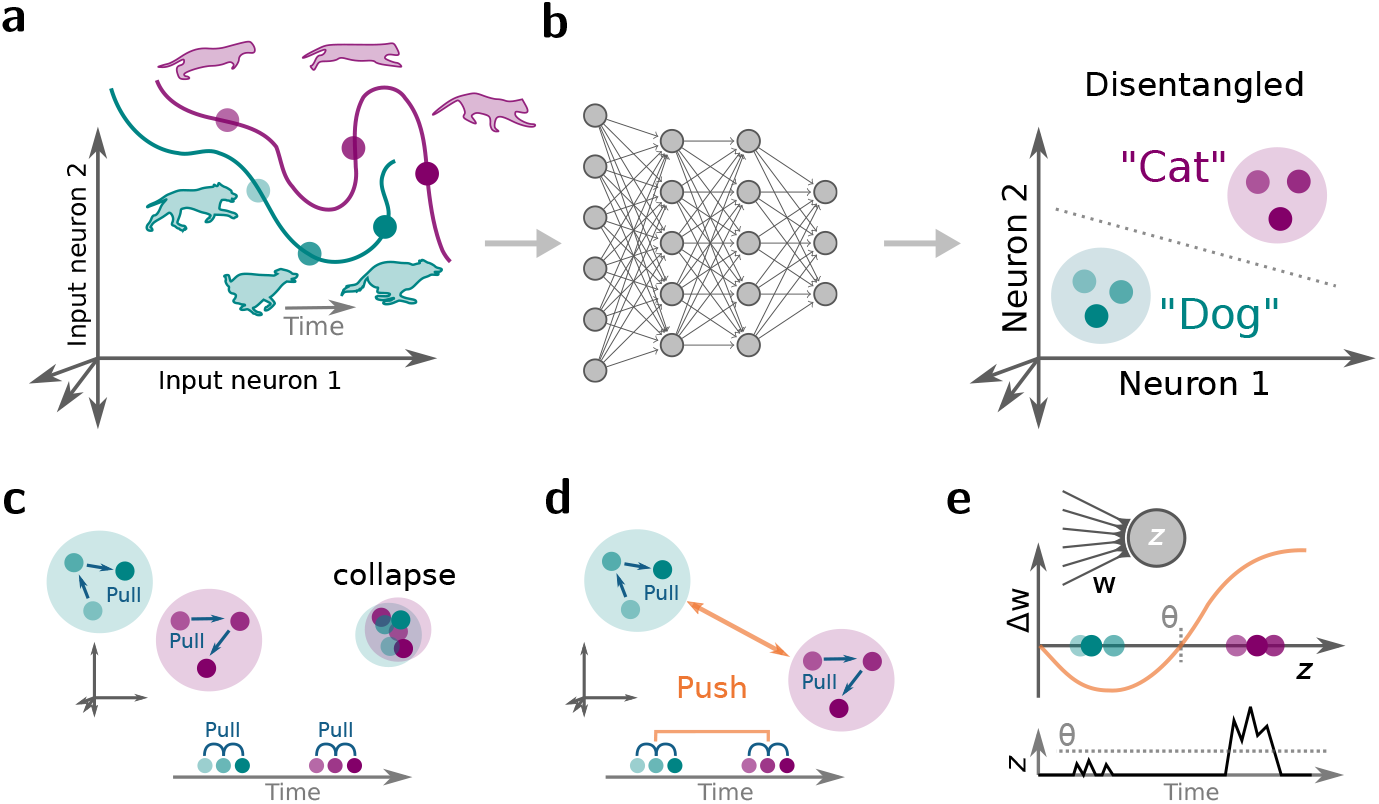
Disentangling sensory stimuli with plastic neural networks. **(a)** Schematic of an evoked response in sensory input neurons. The neuronal response patterns for distinct stimuli correspond to points in a high dimensional space spanned by the neuronal activity levels. The response patterns from different stimulus classes, e.g., cats and dogs, form a low-dimensional manifold in the space of all possible response patterns. Generally, different class manifolds are entangled, which means that the stimulus identity cannot be readily decoded from a linear combination of the neuronal activities. **(b)** Sketch of a deep neural network (left) that transforms inputs into disentangled internal representations that are linearly separable (right). **(c)** Schematic of how predictive learning influences latent representations (left). Learning tries to “pull” together representations that frequently co-occur close in time (bottom). However, without opposing forces, such learning dynamics lead to representational “collapse” whereby all inputs are mapped to the same output and thereby become indistinguishable (right). **(d)** Self-supervised learning (SSL) avoids collapse by adding a repelling force that acts on temporally distant representations that are often semantically unrelated. **(e)** Plot of postsynaptic neuronal activity *z* over time (bottom) and the Bienenstock-Cooper-Munro (BCM) learning rule (top; 7, 8) which characterizes the sign and magnitude of synaptic weight change Δ*w* as a function of postsynaptic activity *z*. Notably, the sign of plasticity depends on whether the evoked responses are above or below the plasticity threshold *θ*. Using the example of Neuron 1 in panel (b), the BCM rule potentiates synapses that are active when a “Cat” stimulus is shown, whereas “Dog” stimuli induce long-term depression (LTD). This effectively pushes the evoked neuronal activity levels corresponding to both stimuli away from one another, and thereby prevents representational collapse.

Unfortunately, standard deep learning methods are difficult to reconcile with biology. On the one hand, they rely on backpropagation, an algorithm considered biologically implausible, although neurobiology may implement effective alternatives [5, 9–12]. On the other hand, humans and animals cannot learn through strong label-based supervision, as this would require knowledge of a label for *every* input pattern.

Here, self-supervised learning (SSL), a family of unsupervised machine learning algorithms, may offer a remedy. SSL does not need labeled data but instead relies on prediction, a notion also supported by neurobiology [13–18]. Prediction can happen in the input space by, for instance, reconstructing one part of an image from another, as for autoencoders [19], or by predicting the next word in a sentence, as done in language models. Alternatively, prediction can occur in latent space by requiring internal representations of related inputs to predict each other [20, 21]. Latent space prediction is more compelling from a neuroscience perspective since it does not require an explicit decoder network that computes prediction errors at the input, i.e., the sensory periphery, for which there is little experimental support. Instead, latent prediction errors are computed locally or at network outputs (cf. Fig. 1) and drive learning by “pulling” together related internal representations for stimuli that frequently occur close in time (Fig. 1c), similar to slow feature analysis (SFA) [22, 23].

However, a major issue with this strategy is that without any forces opposing this representational pull, such learning inevitably leads to “representational collapse,” whereby all inputs are mapped to the same internal activity pattern which precludes linear separability (Fig. 1c). One typical solution to this issue is to add forces that “push” representations corresponding to *different* unrelated stimuli away from one another (Fig. 1d). This is usually done by invoking so-called “negative samples,” which are inputs that do not frequently occur together in time. This approach has been linked to biologically plausible three-factor learning rules [24, 25], but it requires constantly switching the sign of plasticity depending on whether two successive inputs are related to each other or not. Yet, it is unknown whether and how such a rapid sign switch is implemented in the brain.

Another possible solution for avoiding representational collapse without negative samples is to prevent neuronal activity from becoming constant over time, for instance, by maximizing the variance of the activity [26]. Interestingly, variance maximization is a known signature of Hebbian plasticity [27, 28], which has been found ubiquitously in the brain [29, 30]. While Hebbian learning is usually thought of as the primary plasticity mechanism rather than playing a supporting role, Hebbian plasticity alone has had limited success at disentangling representations in DNNs [5, 31, 32].

This article introduces Latent Predictive Learning (LPL), a conceptual learning framework that overcomes this limitation and reconciles SSL with Hebbian plasticity. Specifically, the local learning rules derived within our framework combine a plasticity threshold, as observed in experiments (Fig. 1e; [8, 29, 33–35]), with a predictive component, inspired by SSL and SFA, that renders neurons selective to temporally contiguous features in their inputs. When applied to the layers of deep hierarchical networks, LPL yields disentangled representations of objects present in natural images without requiring labels or negative samples. Crucially, LPL effectively disentangles representations as a local learning rule without requiring explicit spatial credit assignment mechanisms. Still, credit assignment capabilities can further improve its effectiveness. We demonstrate that LPL captures central findings of unsupervised visual learning experiments in monkeys and in spiking neural networks (SNNs) and naturally yields a classic spike-timing-dependent plasticity (STDP) window, including its experimentally observed firing-rate-dependence [29]. These findings suggest that LPL constitutes a plausible normative plasticity mechanism that may underlie representation learning in biological brains.

## Results

To study the interplay of Hebbian and predictive plasticity in sensory representation learning, we derived a plasticity model from an SSL objective function that is reminiscent of and extends the classic BCM learning rule [7, 8] (Methods; Supplementary Note S1). According to our learning rule, the temporal dynamics of a synaptic weight *W_j_* are given by

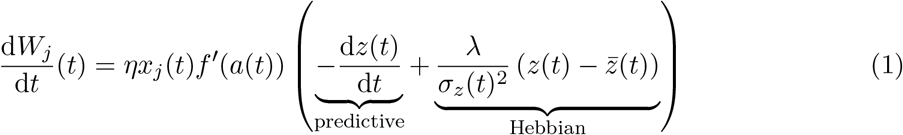

where *η* is a small positive learning rate, *x_j_*(*t*) denotes the activity of the presynaptic neuron *j*, *z*(*t*) = *f*(*a*(*t*)) is the neuronal activity with the activation function *f*, and the net input current *a*(*t*) ∑_*k*_ *W_k_x_k_*(*t*). We call the first term in parentheses the predictive term because it promotes learning of slow features [22, 23] by effectively “pulling together” postsynaptic responses to temporally consecutive input stimuli. Importantly, it cancels when the neural activity does not change and, therefore, accurately *predicts* future activity. In the absence of any additional constraints, the predictive term leads to collapsing neuronal activity levels [22]. In our model, collapse is prevented by the Hebbian term in which 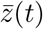, the running average of the neuronal activity appears, which is reminiscent of BCM-theory [7, 8]. Its strength further depends on an online estimate of the postsynaptic variance of neuronal activity 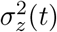. This modification posits an additional metaplasticity mechanism controlling the balance between predictive and Hebbian plasticity depending on the postsynaptic neuron’s past activity.

To make the link to BCM explicit, we rearrange the terms in Eq. (1) to give:

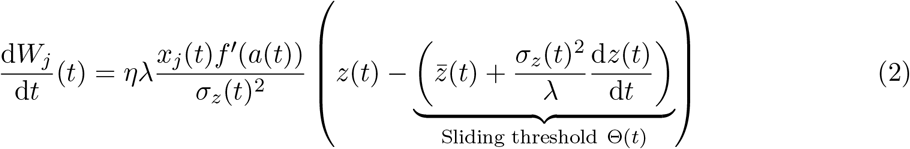

where Θ(*t*) corresponds to a time-dependent sliding plasticity threshold (cf. Fig. 1e). While the precise shape of the learning rule depends on the choice of neuronal activation function, its qualitative behavior remains unchanged as long as the function is monotonic (Extended Data Fig. 1). Despite the commonalities, however, there are three essential differences to the BCM model. First, in our model, the threshold depends only linearly on 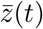 (Extended Data Fig. 1b), whereas in BCM, the threshold is typically a supralinear function of the moving average 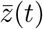. Second, the added dependence on the predictive term 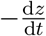 constitutes a separate mechanism that modulates the plasticity threshold depending on the rate-of-change of the postsynaptic activity (Extended Data Fig. 1c,d). Third, our model adds a variance-dependence that has diverse effects on the sliding threshold when the neuronal output does not accurately predict future activity and, thus, changes rapidly (Extended Data Fig. 1c,d). We will see that these modifications are crucial to representation learning from the temporal structure in sensory inputs. Because the predictive term encourages neurons to predict future activity at their output, and thus in latent space rather than the input space, we refer to Eq. (1) as the Latent Predictive Learning (LPL) rule.

### LPL finds contiguous features in temporal data

To investigate the functional advantages of LPL over BCM and other classic Hebbian learning rules (Supplementary Note S2), we designed a synthetic two-dimensional learning task in which we parametrically controlled the proportion of predictable changes between subsequent observations (Fig. 2a; Methods). The data sequence consisted of noisy inputs from two clusters separated along the *x*-axis. Consecutive inputs had a high probability of staying within the same cluster, thus making cluster identity a temporally contiguous feature. By varying the noise amplitude *σ_y_* in the *y*-direction, we controlled the amount of unpredictable changes. We simulated a single rate neuron with different datasets for varying *σ_y_*, while the two input connections were plastic and evolved according to the LPL rule (Eq. (1)) until convergence. We then measured neuronal selectivity to cluster identity (Methods).

**Figure 2:**
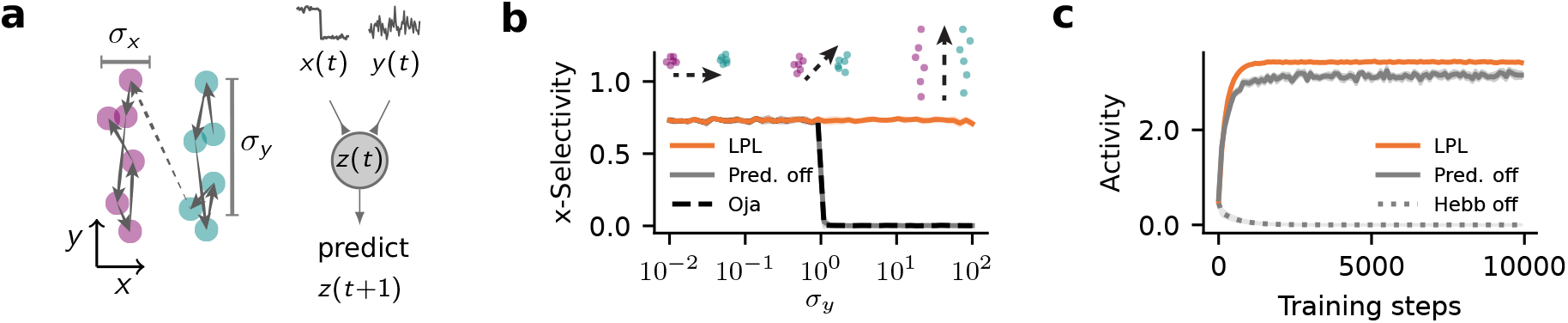
LPL learns predictive features. **(a)** Illustration of the two-dimensional synthetic data generating process. Consecutive data points stay within the same cluster separated along the *x*-direction and are drawn independently from the corresponding normal distribution centered in that cluster (left). These data are fed into a linear neuron that learns via LPL (right). **(b)** Cluster selectivity of the features learned by LPL with and without the predictive term and by Oja’s rule for different values of *σ_y_*. By varying *σ_y_*, we obtain a family of sequences with different amplitudes of within-cluster transitions (top). LPL selects temporally contiguous features and therefore ensures that the neuron always becomes selective to cluster identity. Oja’s rule finds PC1, the direction of highest variance, which switches to the noise direction at *σ_y_* = 1. LPL without the predictive component shows the same behavior. Selectivity values were averaged over ten random seeds. The shaded area corresponds to one SD. **(c)** Mean output activity of the neuron over training time for *σ_y_* = 1 under different versions of LPL. LPL initially increases its response and saturates at some activity level, even when the predictive term is disabled. However, without the Hebbian term, the activity collapses to zero.

We found that LPL rendered the neuron selective to the cluster identity for a large range of *σ_y_* values (Fig. 2b). However, without the predictive term, the selectivity to cluster identity was lost for large *σ_y_* values. This behaviour was expected because omitting the predictive term renders the learning rule purely Hebbian, which biases selectivity toward directions of high variance. To illustrate this point, we repeated the same simulation with Oja’s rule, a classic Hebbian rule that finds the principal component in the input, and found similar qualitative behaviour. Thus LPL behaves fundamentally different from purely Hebbian rules, by selecting predictable features in the input.

To confirm that the Hebbian term is essential for LPL to prevent representational collapse, we simulated learning without the Hebbian term (cf. Eq. (1)). We observed that the neuron’s activity collapses to zero firing rate as expected (Fig. 2c). Conversely, learning with the Hebbian term but without the predictive term did not result in collapse. Therefore, LPL’s Hebbian component is essential to prevent activity collapse.

Moreover, Hebbian plasticity needs to be dynamically regulated to prevent run-away activity [36]. In LPL this regulation is achieved by inversely scaling the Hebbian term by a moving estimate of the variance of the postsynaptic activity 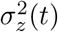. Without this variance-modulation, neural activity either collapsed or succumbed to runaway activity depending on which term was dominant (Supplementary Note S3). Either case precluded the neuron from developing cluster selectivity. We verified that these findings generalized to higher-dimensional tasks with more complex co-variance structure (Supplementary Note S4). Hence, the combination of the predictive with variance-modulated Hebbian metaplasticity in LPL is needed to learn invariant predictive features independently of the co-variance structure in the data.

### LPL disentangles representations in deep hierarchical networks

As we move through the world, we see objects, animals, and people under different angles and contexts (Fig. 3a). Therefore, objects themselves constitute temporally contiguous features in normal vision. We thus wondered whether training an artificial DNN with LPL on image sequences with such object permanence results in disentangled representations. To that end, we built a convolutional DNN model in which we “stacked” layers whose synaptic connections evolved according to the LPL rule. Additionally we included a term to decorrelate neurons within each layer. Inhibitory plasticity presumably plays this role in biological neural networks [37–40]. LPL was implemented in a “layer-local” manner, meaning that there was no backpropagation through layers (Methods).

**Figure 3:**
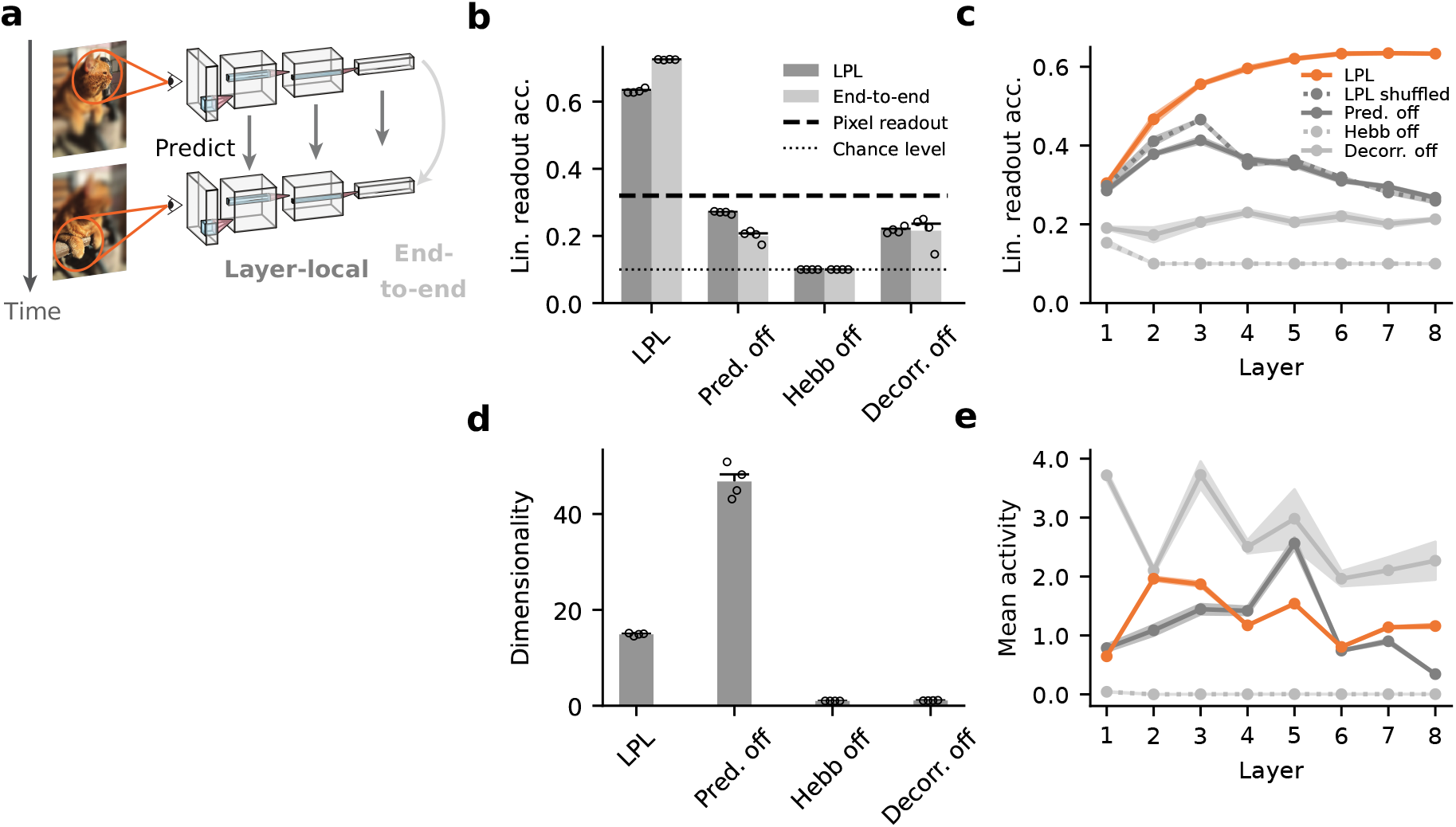
LPL disentangles representations in DNNs. **(a)** Schematic of the DNN trained using LPL. We distinguish two learning strategies: Layer-local and end-to-end learning. In layer-local LPL, each layer’s learning objective 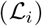 is to predict representations within the same layer, whereas end-to-end training takes into account the output layer representations only 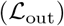, and updates hidden layer weights using backpropagation. **(b)** Linear readout accuracy of object categories decoded from representations at the network output after training it on natural image data (STL-10; see Methods for details) with different learning rules in layer-local (dark) as well as the end-to-end configuration (light). “Pred. off” corresponds to LPL but without the predictive term in the learning rule (cf. Eq. 7). “Hebb off” refers to the configuration without the BCM-like Hebbian term. Finally, “Decorr. off” is the same as the single neuron learning rule (Eq. (1)) without the decorrelation term. LPL yields features with high linear readout accuracy. In contrast, when any component of LPL is disabled, linear readout accuracy drops below the pixel-decoding accuracy of ~32% (dashed line). Error bars indicate standard error of the mean (SEM) over four trials. **(c)** Linear readout accuracy of the internal representations at different layers of the DNN after layer-local training. LPL’s representations improve up to six layers and then settle at a high level. In contrast, readout accuracy is close to chance level without the Hebbian component, and similarly remains at low levels when the decorrelating mechanism is switched off. Interestingly, when the predictive term is off, the readout accuracy initially increases in early layers, but then ultimately decreases back below the pixel-level accuracy with further increasing depth. Finally, the full LPL learning rule applied to inputs in which temporal contingency is destroyed (LPL shuffled) behaves qualitatively similar to the purely Hebbian rule. **(d)** Dimensionality of the internal representations for the different learning rule configurations shown in (b). When either the Hebbian or decorrelation term are disabled, the dimensionality of the representations collapses to one. **(e)** Mean neuronal activity at different layers of the DNN after training with the different learning rule variants shown in (c). Excluding the Hebbian term (dotted line) leads to collapsed representations in all layers.

To simulate temporal sequences of related visual inputs, we generated pairs of images sampled from a large dataset, by applying different randomized transformations (Extended Data Fig. 2; Methods). We trained our network model on these visual data until learning converged and evaluated the linear decodability of object categories from the learned representations using a separately trained linear classifier.

We found that in networks trained with LPL, object categories could be linearly decoded at the output with an accuracy of (63.2 ±0.3)% (Fig. 3b; Table 1), suggesting that the network has formed partially disentangled representations (Extended Data Fig. 3). To elucidate the roles of the different learning rule components, we conducted several ablation experiments. First, we repeated the same simulation but now excluding the predictive term. This modification resulted in an accuracy of (27.0 ± 0.2)%, which is *lower* than the linear readout accuracy of a classifier trained directly on the pixels of the input images (see Table 1), indicating that the network did not learn disentangled representations, consistent with previous studies on purely Hebbian plasticity [5, 32]. We measured a similar drop in accuracy when we disabled either the Hebbian or the decorrelation component during learning (Fig. 3b).

**Table 1:** Linear classification accuracy on the STL-10 and CIFAR-10 datasets for LPL and for a linear decoder trained on the raw pixel values (Methods). Error values correspond to SEM over four simulations with different random seeds.

|  | STL-10 |  | CIFAR-10 |  |
| --- | --- | --- | --- | --- |
|  | Layer-local (%) | End-to-end (%) | Layer-local (%) | End-to-end (%) |
| DNN with LPL | 63.2±0.3 | 72.5±0.1 | 59.4±0.4 | 70.4±0.2 |
| Raw pixel values | 31.6 |  | 35.9 |  |

Convolutional DNNs trained through supervised learning use depth to progressively separate representations [2]. To understand whether networks trained with LPL similarly leverage depth, we measured the linear readout accuracy of the internal representations at every layer in the network. Crucially, we found that in the LPL-trained networks, the readout accuracy increased with the number of layers until it gradually saturated (Fig. 3c), whereas this was not the case when any of the components of LPL was disabled. Similarly, readout accuracy decreased, when the temporal contiguity in the input was broken by shuffling, reminiscent of experiments in developing rats [17]. Together, these results suggest that LPL’s combination of Hebbian, predictive, and decorrelating elements is crucial for disentangling representations in hierarchical DNNs.

In SSL, the two most common causes for failure to disentangle representations are representational and dimensional collapse (Extended Data Fig. 4), due to excessively high neuronal correlations [41]. To disambiguate between these two possibilities in our model, we computed the dimensionality of the representations and the mean neuronal activity at every layer (Methods). We found that disabling either the Hebbian or the decorrelation component led to a dimensionality of approximately one, whereas the LPL rule with and without the predictive term resulted in higher dimensionality: ≈ 15 or ≈ 50 respectively (Fig. 3d). Disabling the Hebbian term silenced all layers (Fig. 3e), demonstrating representational collapse. In contrast, disabling the decorrelation term resulted in non-zero activity levels, indicating that dimensional collapse underlies its poor readout accuracy (Fig. 3e). Finally, we verified that excluding LPL’s predictive component neither caused representational nor dimensional collapse, suggesting that the decreasing linear readout accuracy with depth was due to the network’s inability to learn good internal representations. Taken together, these results show that the predictive term is crucial for disentangling object representations in DNNs (Fig. 3) whereas the other terms are essential to prevent different forms of collapse.

It is an ongoing debate whether neurobiology implements some form of credit assignment [5, 9–12]. Above we showed that LPL, as a local learning rule, effectively disentangles representations without the need for credit assignment, provided mechanisms exist that ensure neuronal decorrelation [38]. Naturally, our next question was whether a non-local LPL formulation could improve learning. To that end, we considered the fully non-local case using backpropagation. Specifically, we repeated our simulations with end-to-end training on the LPL objective defined at the network’s output (Methods). While we do not know how the brain would implement such a non-local LPL algorithm, it provides an upper performance estimate of what is possible. End-to-end learning reproduced all essential findings of layer-local learning while increasing overall performance (Fig.3b; Table 1). Thus LPL’s performance improves in the non-local setting, further underscoring that biological networks could benefit from credit assignment circuit mechanisms.

The above simulations used pairs of augmented images. To check whether the key findings generalized to more realistic input paradigms and other measures of disentangling, we trained DNNs with LPL on procedurally generated videos from the 3D Shapes dataset [42]. The videos consisted of objects shown under a slowly changing view angle, scale, or hue and occasional discontinuous scene changes, but without additional image augmentation (Extended Data Fig. 5a,b; Methods). We found that LPL-trained networks reliably disentangle object identity. In contrast, networks trained without predictive learning failed to do so (Extended Data Fig. 5c). Finally, the ground-truth latent manifold structure in the procedurally generated dataset is known. This knowledge allowed us to probe disentangling of the latent manifold directly instead of using linear classification as a proxy. This analysis revealed that LPL-trained networks faithfully disentangled the underlying objects and factors. At the same time, they also learned the topology of the data-generating manifold from the temporal sequence structure (Extended Data Figs. 5d–g and 6). Thus LPL’s ability to disentangle representations generalizes to video stimuli and other measures of disentanglement.

### LPL captures invariance learning in the primate inferotemporal cortex

Changing the temporal contiguity structure of visual stimuli induces neuronal selectivity changes in primate inferotemporal cortex (IT), an unsupervised learning effect described by Li and DiCarlo [14]. In their experiment, a macaque freely viewed a blank screen, with objects appearing in the peripheral visual field at one of two alternative locations relative to the (tracked) center of its gaze, prompting the macaque to perform a saccade to this location (Fig. 4a). The experimenters differentiated between normal exposures in which the object does not change during the saccade and “swap exposures” in which the initially presented object was consistently swapped out for a different one as the monkey saccaded to a specific target location X_swap_. Hence, swap exposures created an “incorrect” temporal association between one object at position X_swap_ and a different one at the animal’s center of gaze X_c_. For any particular pair of swap objects, either the location above or below the center of gaze was chosen as X_swap_, and transitions from the opposite peripheral position X_nonswap_ to the center X_c_ were kept consistent as a control. The authors found systematic position- and object-specific changes of neuronal selectivity due to swap exposures which they attributed to unsupervised learning. Specifically, a neuron initially selective to an object *P* over another object *N*, reduced or even reversed its selectivity at the swap position X_swap_ while preserving its selectivity at the non-swap position X_nonswap_ (Fig. 4b).

**Figure 4:**
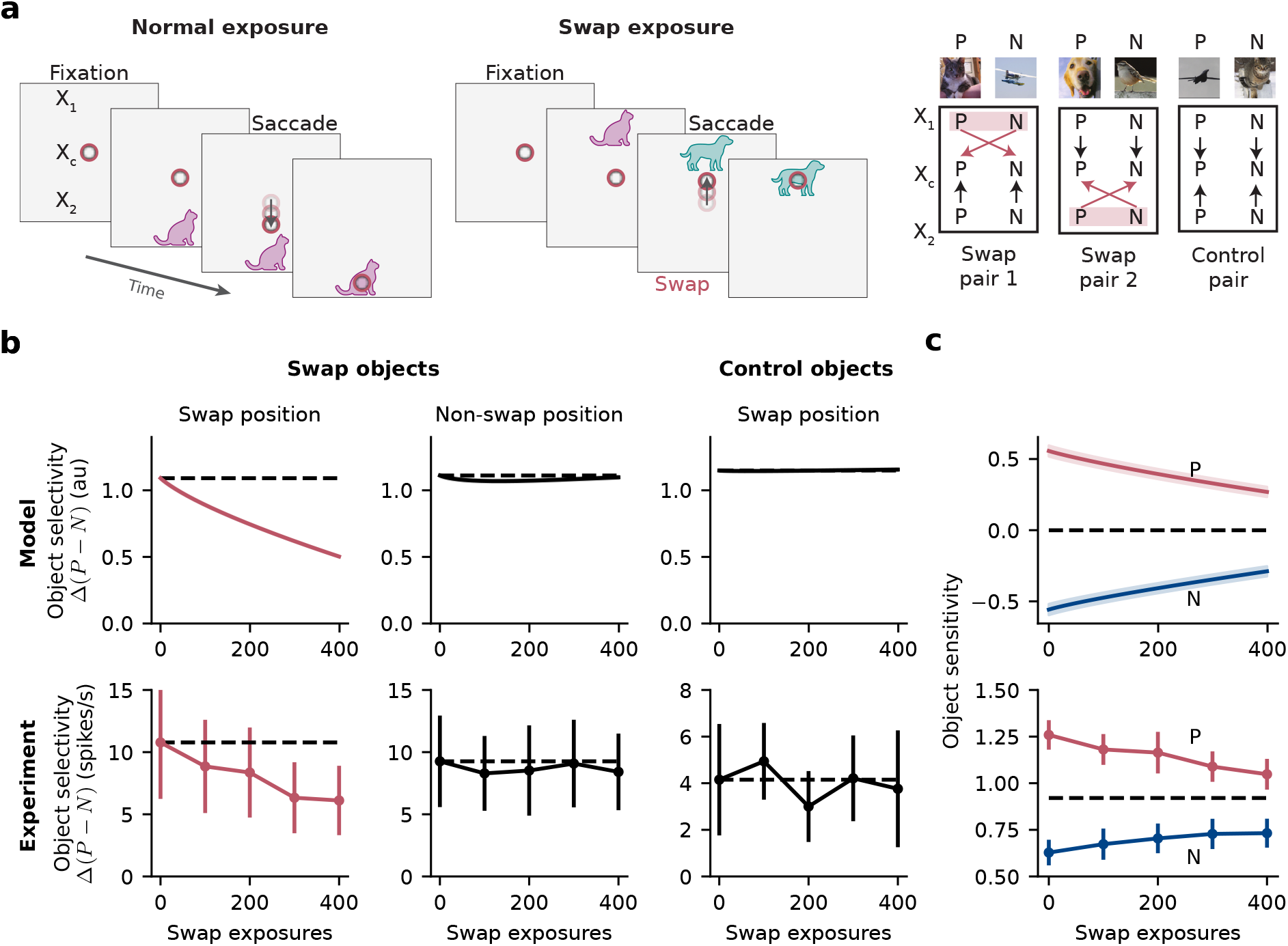
LPL captures invariance learning in the primate inferotemporal cortex (IT). **(a)** Schematic of the simulation setup modeled after the experiment by Li and DiCarlo [14]. The inputs to the model consist of images of objects presented at three different positions X_1_, X_c_, and X_2_ on a blank canvas. Following the original experiment, we performed a targeted perturbation in the simulated visual experience that the model network was exposed to (left and center). Specifically, we switched object identities during transitions from a specific peripheral position, say X_1_, to the central position X_c_, while keeping transitions from the other peripheral position to the center unmodified (right). **(b)** Evolution of object selectivity as a function of number of swap exposures in the model (top row) and observed in-vivo (bottom row; data points extracted and replotted from [14], see Methods for details). We differentiate between pairs of swapped objects at the Swap (left) and Non-swap positions (center) as well as control objects at the Swap position (right). LPL qualitatively reproduces the evolution of swap position-specific remapping of object selectivity as observed in IT. Control objects at the Swap position, i.e., images not used during the swap training protocol, show no selectivity changes in agreement with the experiment. **(c)** Average response to objects *P* and *N* as a function of number of swap exposures. The change in object selectivity between preferred objects *P* and non-preferred objects *N* is due to both increased responses to *N* and decreased responses to *P* in both our model (top) and the experimental recordings (bottom).

We wanted to know whether LPL can account for these observations. To that end, we built a DNN model and generated input images by placing visual stimuli on a larger gray canvas to mimic central and peripheral vision as needed for the experiment (cf. Fig. 4a; Methods). Importantly, we ensured that the network’s input dimension and output feature map size were large enough to avoid full translation invariance due to the network’s convolutional structure alone. To simulate the animal’s prior visual experience, we trained our network model with LPL on a natural image dataset. After training, the learned representations were invariant to object location on the canvas (Extended Data Fig. 7), a known property of neural representations in the primate IT [1]. Next, we simulated targeted input perturbations analogous to the original experiment. For a given pair of images from different classes, we switched object identities during transitions from a specific peripheral position, say X_1_, to the center X_c_ while keeping transitions from the other peripheral position X_2_ to the center unmodified. We used X_1_ as the swap position for half of the image pairs and X_2_ for the other half. Throughout, we recorded neuronal responses in the network’s output layer while the weights in the network model evolved according to the LPL rule.

We observed that the neuronal selectivity between preferred inputs *P*, as defined by their initial preference (Methods), in comparison to non-preferred stimuli *N* in the model qualitatively reproduced the results of the experiment (Fig. 4b). Effectively, LPL trained the network’s output neurons to reduce their selectivity to their preferred inputs *P* at the swap position while preserving their selectivity at the non-swap position. Furthermore, we observed that object selectivity between pairs of control objects did not change, consistent with the experiment (Fig. 4b). Further analysis revealed that the origin of the selectivity changes between *P* and *N* stimuli at the swap position was due to both increases in responses to *N* and decreases in responses to *P*, an effect also observed in the experiments (Fig. 4c). Thus, LPL can account for neuronal selectivity changes observed in monkey IT during in-vivo unsupervised visual learning experiments.

### Spiking neural networks with LPL selectively encode predictive inputs

So far we considered LPL in discrete-time rate-based neuron models without an explicit separation of excitatory and inhibitory neurons. In contrast, cortical circuits consist of spiking neurons that obey Dale’s law, and learn in continuous time. To test whether our theory would extend to such a more realistic setting, we simulated a plastic recurrent SNN model consisting of 100 excitatory and 25 inhibitory sparsely connected neurons (Fig. 5a; Methods). The circuit received input from five Poisson populations of 100 neurons each, whose temporally varying firing rates encoded signals with different temporal properties (Fig. 5b; Methods). Input population *P*0 had a constant firing rate, whereas *P*1’s and *P*2’s firing rates followed two independent slowly varying signals. We also defined two control populations *P*1_ctl_ and *P*2_ctl_ whose firing rates were temporally shuffled versions of *P*1 and *P*2 using bins of 10 ms duration. Crucially, all populations had the same mean firing rate of 5 Hz. The input connections to the excitatory neurons evolved according to the spiking LPL rule (cf. (1)), a local learning rule without the decorrelation term. Decorrelation was achieved through inhibitory STDP (38; Methods).

**Figure 5:**
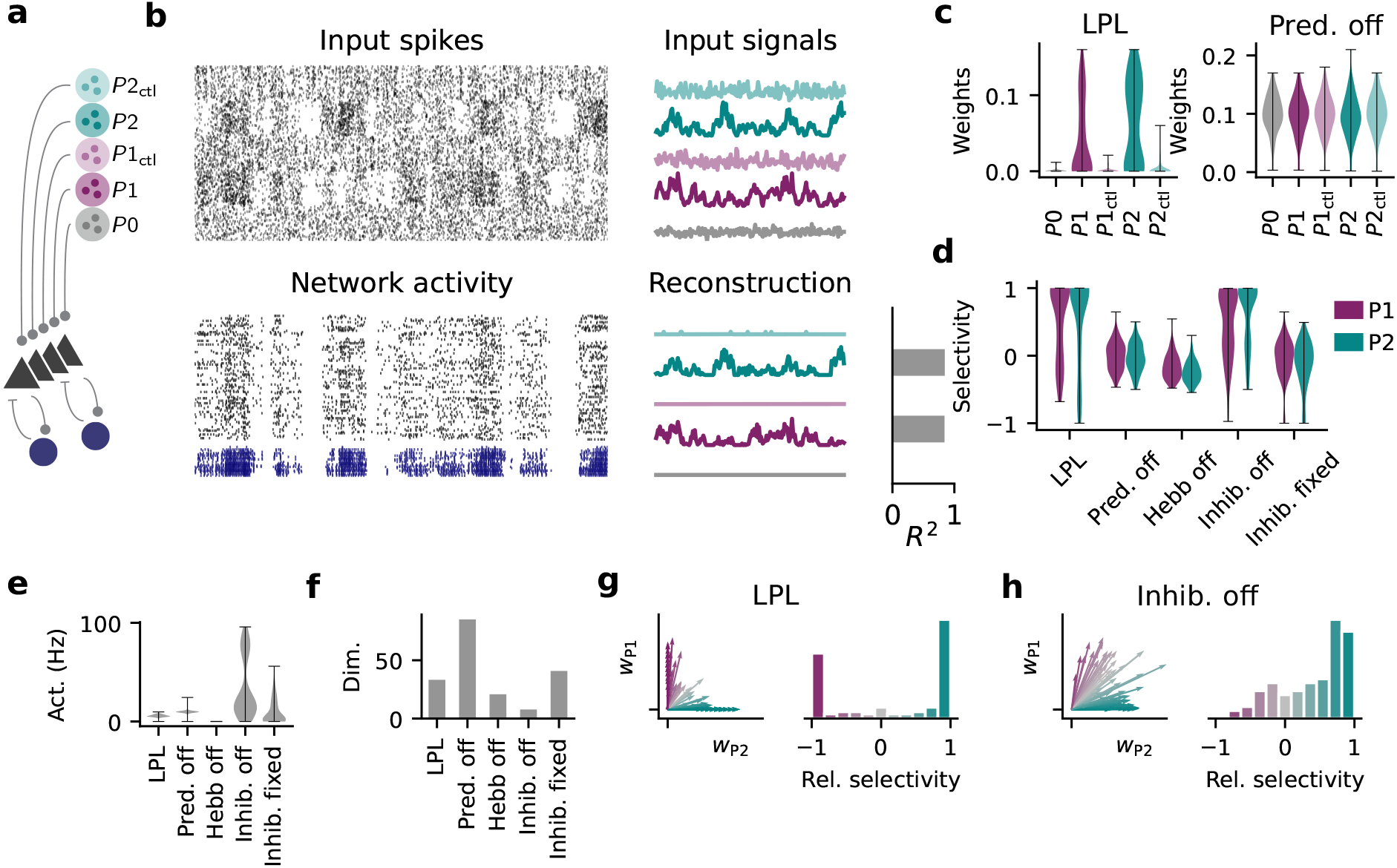
LPL in a spiking neural network (SNN). **(a)** Wiring diagram of the SNN with five distinct input populations. **(b)** Snapshot of spiking activity over 100 ms after LPL plasticity for the inputs (top left) and the network (bottom left) separated into excitatory (black) and inhibitory neurons (blue). The input spikes are organized in five distinct Poisson populations whose firing rates evolve according to five different temporal input signals (top right). Population activity of two slowly varying signals (*P*_1/2_) can be linearly reconstructed (Methods) with high *R*^2^ values from the network activity whereas temporally shuffled control signals (“ctl”; Methods) are heavily suppressed (bottom right). **(c)** Distribution of synaptic connection strengths grouped by input population. Input connections from slowly varying signals are larger than those from the shuffle controls (left), but not when learning with the predictive term turned off (right). **(d)** Signal selectivity as relative difference between signal and control pathway for networks trained with different learning rule variations (Methods). “LPL” refers to learning with the spiking LPL rule combined with inhibitory plasticity on the inhibitory-to-excitatory connections. “Pred. off” corresponds to learning without the predictive term, and “Hebb off” to learning without the Hebbian term. “Inhib. off” refers to a setting without any inhibitory neurons, whereas “Inhib. fixed” indicates a setting where the inhibitory-to-excitatory weights are held fixed. The network with LPL and inhibitory plasticity acquires high selectivity to both signals. Selectivity is lost if the predictive term, the Hebbian term, or inhibitory plasticity are switched off. When inhibition is removed altogether, selectivity remains but is significantly reduced. Error bars indicate SEM over all excitatory neurons. **(e)** Average firing rate of excitatory neurons in the network for the different configurations in (d). When the Hebbian term is off, spiking activity collapses to low activity levels in contrast to all other configurations in which it settles at intermediate activity levels. **(f)** Dimensionality of the neuronal representations (Methods) for the different configurations in (d). Inhibition prevents dimensionality collapse, even in cases where inhibition is not plastic. **(g)** Averaged weight vectors of all excitatory neurons corresponding to input populations *P*1 and *P*2 (left) and the distribution of relative neuronal selectivities between these populations (right). Most neurons become selective either to *P*1 or *P*2, but few to both signals simultaneously. Color indicates relative preference of their weight vectors to either signal (Methods). **(h)** Same as **(g)**, but without an inhibitory population. Most neurons develop selectivity to *P*2 or mixed selectivity to both signals, and their weight vectors are more correlated.

We ran the SNN model for approximately 28 h of simulated time, at which point the network’s firing dynamics had settled into an asynchronous irregular activity regime from which the slowly varying input signals could be decoded linearly with high fidelity (Fig. 5b). In contrast, the rate fluctuations of the shuffled control signals (*P*1_ctl_ and *P*2_ctl_) and the constant firing-rate input (*P*0) could not be reconstructed linearly with high accuracy, consistent with the idea that the network preferentially represents the slowly varying inputs in its activity. This notion was further supported by computing the mean connection strength of the afferent connectivity matrix (Fig. 5c). We further computed the relative difference between the average afferent weight from each signal in comparison to its associated control pathway. As expected, we found that neuronal weights were preferentially tuned to the slow input channels (Fig. 5d). However, this selectivity was lost when we turned either the predictive or the Hebbian term off. The absence of Hebbian plasticity was further accompanied by activity collapse (Fig. 5e), like in the rate-based network.

To investigate the role of inhibition in successful representation learning in the SNN, we repeated the above simulation without the inhibitory population. This manipulation resulted in excessively high firing rates (Fig. 5e; Extended Data Fig. 8), a notable reduction of the representational dimensionality (Fig. 5f; Methods), and lower selectivity to the slow signals (Fig. 5d). The reasons for this reduction can be seen in the distribution of weight vectors. In the network with plastic inhibition, weights were more decorrelated and exclusively selective to either *P*1 or *P*2 (Fig. 5g). In contrast, removing inhibition resulted in more correlated weights with few neurons preferentially tuned to one signal or the other (Fig. 5h). Finally, we repeated the same simulation in a network with inhibitory neurons, but without inhibitory plasticity. This manipulation led to comparable representational dimensionality as for LPL (Fig. 5f), but caused a loss of selectivity relative to the shuffled controls (Fig. 5d). These results indicate that inhibition is needed to prevent correlated neuronal activity and the ensuing reduction in representational dimensionality. Further, inhibitory plasticity is required to ensure that the slow signals are preferentially represented (Extended Data Fig. 8). Together, these findings illustrate that LPL learns predictive features in realistic spiking circuits with separate excitatory and inhibitory neuronal populations.

### LPL qualitatively reproduces experimentally observed rate and spike-timing dependence of synaptic plasticity

Next, we wanted to examine whether the spike-based LPL rule is consistent with experimental observations of plasticity induction. Experiments commonly report intertwined rate and spike-timing dependence presumably mediated through nonlinear voltage- and calcium-dependent cellular mechanisms [30, 43]. Theoretical work has further established conceptual links between phenomenological STDP models, SFA, and BCM theory [23, 44–48].

To compare LPL to experiments, we simulated a standard STDP induction protocol. Specifically, we paired 100 pre- and post-synaptic action potentials with varying relative timing Δ*t* for a range of different repetition frequencies *ρ*. During the entire plasticity induction protocol, the postsynaptic cell was kept depolarized close to its firing threshold and weights evolved according to spike-based LPL. We repeated the simulated induction protocol for different initial values of the slowly moving averages of the postsynaptic firing rate 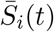 and variance 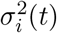 (Methods). This was done because these variables do not change much over the course of a single induction protocol due to their slow dynamics. Their presence, however, makes LPL a form of metaplasticity, i.e., plasticity depends on past neuronal activity.

We found that for small initial values of 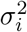, the induced weight changes followed an antisymmetric temporal profile consistent with STDP experiments (Fig. 6a). For larger initial values of 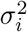, the STDP window changed to a more symmetric and then ultimately an anti-Hebbian profile while the plasticity amplitude was suppressed, as expected due to the variance-dependent suppression of the Hebbian term in the learning rule (Fig. 6b,c). Next we investigated the effect of different initial values for 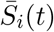, which acts as a moving threshold reminiscent of BCM. Specifically, we recorded plastic changes at two fixed spike timing intervals Δ*t* = ±10 ms for 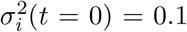. For intermediate threshold values 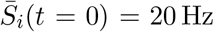, causal spike timing induced long-term potentiation (LTP) with a nonlinear frequency dependence (Fig. 6d) whereas acausal pre-after-post timings showed a characteristic cross-over from LTD to LTP similarly observed in experiments [29]. In contrast, a low initial threshold 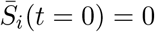, which would occur in circuits that have been quiescent for extended periods of time, resulted in LTP induction for both positive and negative spike timings whereas a high initial value 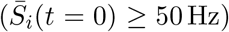, corresponding to circuits with excessively high activity levels, led to LTD (Extended Data Fig. 9). Importantly such slow shifts in activity-dependent plasticity behavior are consistent with the metaplasticity observed in monocular deprivation experiments [8, 34, 48]. Thus, LPL qualitatively captures key phenomena observed in experiments such as STDP, the rate-dependence of plasticity, and metaplasticity, despite not being optimized to reproduce these phenomena. Rather our model offers a simple normative explanation for the necessity of different plasticity patterns that are also observed experimentally [43].

**Figure 6:**
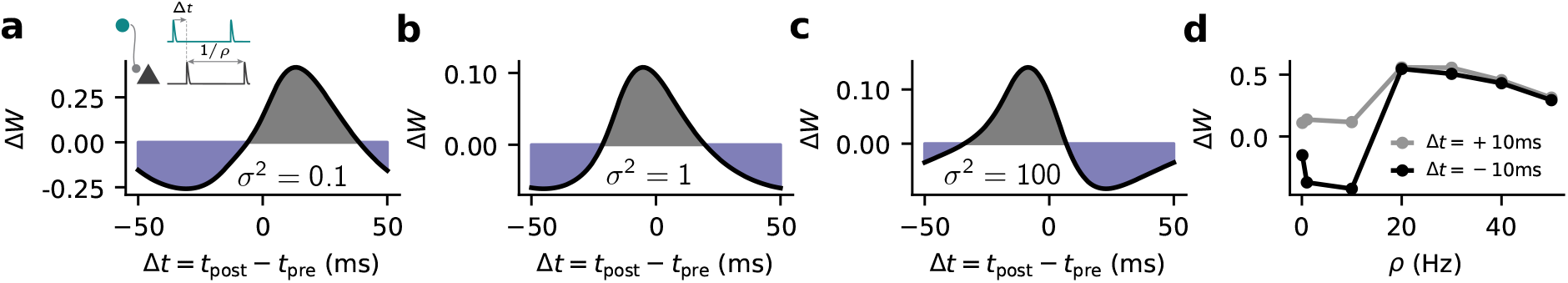
LPL accounts for STDP and predicts metaplasticity of the STDP window. **(a)** Rela-tive weight change due to LPL in response to a standard STDP induction protocol with varying spike timing Δ*t* for 100 pairings at a repetition frequency of *ρ* = 10 Hz (inset) for an initial value of *σ*^2^(*t* = 0) = 0.1. **(b)** Same as (a), but with initial value of *σ*^2^(0) = 1. **(c)** Same as (a), but with *σ*^2^(0) = 100. **(d)** Relative weight change as a function of repetition frequency *ρ* for positive and negative relative spike timings (Δ*t* = ±10 ms).

## Discussion

In this article, we have introduced LPL, a local plasticity rule that extends BCM theory by adding a predictive component to Hebbian learning. We demonstrated that LPL disentangles latent object representations in DNNs through mere exposure to temporal data in which object identity is preserved across successive inputs provided neuronal activity is decorrelated. Crucially, we show that both predictive and Hebbian learning have to work in symphony to accomplish this. Moreover, we demonstrated that LPL qualitatively captures the representational changes observed in unsupervised learning experiments in monkey IT [14]. Finally, we extended LPL to SNNs and found that the resulting learning rule naturally reproduces STDP and its experimentally observed rate-dependence, while further predicting a new form of metaplasticity with distinct variance-dependent changes to the STDP window.

The idea that sensory networks use temporal prediction as a learning objective to form disentangled internal representations has been studied extensively in both machine learning and neuroscience. The model introduced in this article combines and extends aspects of biologically plausible plasticity models closely related to BCM theory with central ideas from SFA and more recent SSL approaches in machine learning. While SSL has shown great promise in representation learning without labelled data, it is typically formulated as a contrastive learning problem requiring negative samples [20, 21] to prevent representional collapse. However, negative samples explicitly break temporal contiguity during learning and are thus not biologically plausible. LPL does not require negative samples. Instead, it relies on variance regularization as proposed previously to prevent collapse [26]. Our model uses virtually the same mechanism, albeit with a logarithmic variance dependence (Supplementary Note S3), and builds a conceptual bridge from variance regularization to Hebbian metaplasticity.

Like most SSL approaches, Bardes *et al.* [26] used end-to-end learning whereby the objective function is formulated on the embeddings at the network’s output. In contrast, we studied the case of greedy layer-wise learning in which the objective is applied to each layer individually. Doing so alleviates the need for backpropagation and permitted us to formulate the weight updates as local learning rules, similar to work that combined contrastive objectives with greedy layer-wise training [31]. Furthermore, recent work showed that greedy layer-wise contrastive learning is directly linked to plasticity rules that rapidly switch between Hebbian and anti-Hebbian learning through a global third factor [24]. However, both these models required implausible negative samples, whereas LPL does neither require end-to-end training nor negative samples.

LPL shares the shape of the BCM rule, which has been qualitatively confirmed in numerous experimental studies both in-vitro [8, 29, 34] and in-vivo [35]. Furthermore, BCM has been linked to STDP [30] and informed numerous phenomenological plasticity models [44–47, 49]. However, unequivocal evidence for the predicted supralinear behavior of the firing rate-dependence of the BCM sliding threshold remains scarce [34] and the fast sliding threshold required for network stability seems at odds with experiments [36, 48]. In contrast, LPL does not require a rapid nonlinear sliding threshold for stability. Instead, it posits a fast-acting variance-dependence that ensures stability by suppressing Hebbian plasticity when the variance of the output activity is too high. This suppressive effect allows the sliding threshold, which could be implemented by either neuronal or circuit mechanisms [34, 50], to catch up slowly, more consistent with experiments [48]. Hence, LPL offers a possible explanation for the current gap between theory and experiment, while suggesting future investigations of plasticity regulation in neuronal circuits.

The notion of slowness learning has been studied extensively in the context of the Trace Rule [51] and SFA [22, 40] which have conceptual ties to STDP [23]. However, the former enforces a hard constraint on the norm of the weight vector to prevent collapse, while SFA enforces a hard variance constraint. In contrast, LPL implements a soft variance constraint [26] to the same effect. A similar soft constraint on the variance can be derived from statistical independence arguments [52] within a mutual information view of SSL [20]. However, these studies used negative samples, assume rapid global sign switching of the learning rule, and did not connect their work to biological plasticity mechanisms.

Our study has several limitations which we aim to address in future work. First, our study is limited to visual tasks of core object recognition, whereas other sensory modalities may use LPL as a mechanism to form disentangled representations of the external world. For computational feasibility, we restricted ourselves to artificial data augmentation techniques borrowed from SSL and procedurally generated videos with a simple structure, which are only crude proxies of rich real-world stimuli. Finally, there remains a performance gap in classification performance compared to less plausible fully supervised and contrastive approaches (Supplementary Table S1) showing that there remains room for improvement, possibly by incorporating biological circuit mechanisms and top-down feedback connections into the model. It is left as future work to show how LPL can be extended to the circuit level and to more ethologically realistic sensory modalities [53] and video input while further combining them with plausible models of saccadic eye movement.

Despite the limitations, our model makes several concrete predictions about synaptic plasticity. As we have shown, modulating the strength of Hebbian plasticity as a function of the variance of the postsynaptic activity is essential to LPL. A direct prediction of our model is, therefore, that the predictive contribution to plasticity should be best observable when post-synaptic activity is highly variable, while it should be barely observable at low variance levels. While our model does not make quantitative predictions about the time scale on which each neuron would have to estimate its output variance, one would expect that a neuron that has been inactive for a long time, as may be the case in slice experiments, would show stronger Hebbian learning than neurons participating in in-vivo activity. Moreover, LPL should manifest in metaplasticity experiments as a transition from an asymmetric Hebbian STDP window, via a symmetric window to, to ultimately an anti-Hebbian window (cf. Fig. 6) when priming the postsynaptic neuron with increasing output variance. Specifically, we expect a neuron which has remained quiescent for a long period of time to display a classic STDP window, whereas a neuron whose activity has undergone substantial fluctuations in the recent past should show an inverted STDP window. Such metaplasticity may account for the diversity of different shapes of STDP windows observed in experiments [43].

To fathom how established data-driven plasticity models are related to theoretically motivated learning paradigms such as SFA and SSL is essential to understanding the brain. A central open question in neuroscience remains: How do the different components of such learning rules interact with the rich local microcircuitry to yield useful representations at the network level? In this article, we have only scratched the surface by proposing a local plasticity rule and illustrating its aptitude for disentangling internal representations. However, a performance gap remains compared to learning algorithms that can leverage top-down feedback. We expect that extending predictive learning to the circuit and network level will narrow this gap and generate deep mechanistic insights into the underlying principles of neural plasticity.

## Online Methods

### Plasticity model

The LPL rule is derived from an objective function approach. It consists of three distinct parts, each stemming from a different additive term in the following combined objective function:

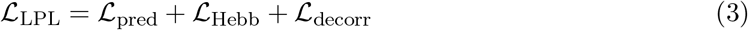

First, the predictive component 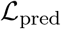 minimizes neuronal output fluctuations for inputs that occur close in time. Second, a Hebbian component 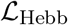 maximizes variance and thereby prevents representational collapse. Finally, 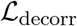 is a decorrelation term that we use in all non-spiking network simulations to prevent excessive correlations between neurons within the same layer in a network. In SNNs decorrelation is achieved without this term through lateral inhibition and inhibitory plasticity.

In the following, we consider a network layer with *N* input units and *M* output units trained on batches of *B* pairs of consecutive stimuli. In all simulations we approximate the temporal derivative ^d*z*^/d*t* which appears in Eqn. (1) by finite differences *z*(*t*) – *z*(*t* –Δ*t*) assuming a discrete timestep Δ*t* while absorbing all constants into the learning rate. In this formulation, the LPL rule has a time horizon of two time steps in the sense that only one temporal transition enters into the learning rule directly. We used this insight to efficiently train our models using minibatches of paired consecutive input stimuli which approximates learning on extended temporal sequences consisting of many time steps. Let *x^b^*(*t*) ∈ ℝ^*N*^ be the input to the network at time *t*, *W* ∈ ℝ^*M*×*N*^ be the weight matrix to be learned, *a^b^*(*t*) = *Wx^b^*(*t*) ∈ ℝ^*M*^ be the pre-activations, and 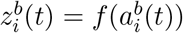 the activity of the *i*th output neuron at time *t*. Finally, *b* indexes the training example within a minibatch of size *B*.

#### Predictive component

We define the predictive objective 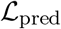 as the mean squared difference between neuronal activity in consecutive time steps.

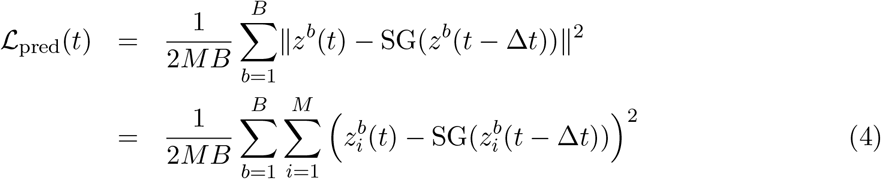

Here SG denotes the Stopgrad function, which signifies that the gradient is not evaluated with respect to quantities in the past.

#### Hebbian component

To avoid representational collapse we rely on the Hebbian plasticity rule that results from minimizing the negative logarithm of the variance of neuronal activity:

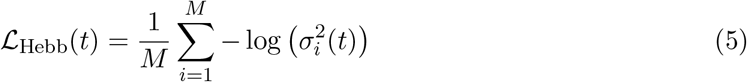

where 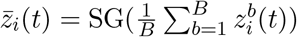 and 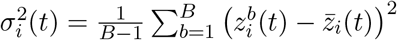 are the current estimates of the mean and variance of the activity of the ith output neuron. Note that we do not compute gradients with respect to the mean estimate, which would require backpropagation through time. Assuming that the mean is fixed allows formulating LPL as a temporally local learning rule (cf. Eq. (3)). To minimize the computational burden in DNN simulations, we performed all necessary computations on minibatches, which includes estimating the mean and variance. However, these quantities could also be estimated using stale estimates from previous inputs, a requirement for implementing LPL as an online learning rule. Using stale mean and variance estimates from previous minibatches in our DNN simulations did cause a drop in readout performance (Supplementary Table S2). Still, such a drop could possibly be avoided either using larger mini batch sizes, by further reducing the learning rate, or by computing the estimates as running averages over past inputs. All of the above manipulations result in essentially the same learning rule (see Supplementary Note S1).

#### Decorrelation component

Finally, we use a decorrelation objective to prevent excessive correlation between different neurons in the same layer as suggested previously [26, 37, 54]. The decorrelation loss function is the sum of the squared off-diagonal terms of the covariance matrix between units within the same layer, which is given as

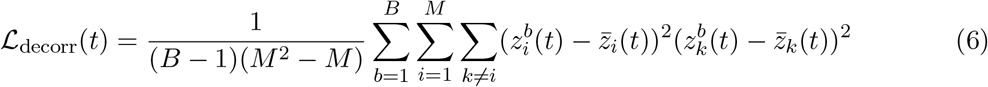

with a scaling factor that keeps the objective invariant to the number of units in the population.

#### The full learning rule

We obtain the LPL rule as the negative gradient of the total objective 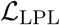 plus an added weight decay. For a single network layer, this yields the layer-local LPL rule where we omitted the time argument t from all present quantities for brevity:

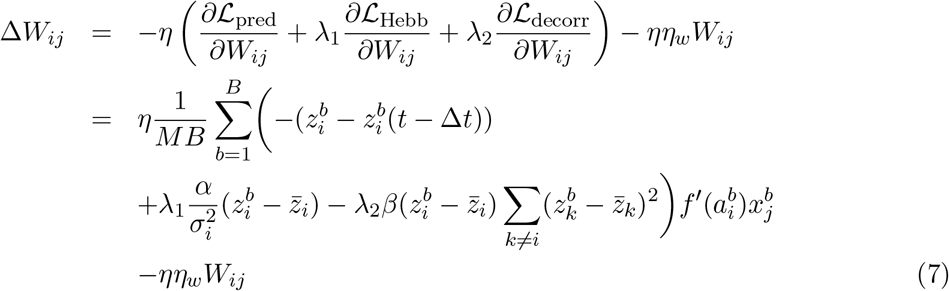

Here λ_1_ and λ_2_ are parameters which control the relative strengths of each objective, *α* and *β* are the appropriate normalizing constants for batch size and number of units, and *η_w_* is a parameter controlling the strength of the weight decay.

#### Numerical optimization methods

We implemented all network models learning with LPL using gradient descent on the equivalent objective function in PyTorch with the Lightning framework. DNN simulations were run on five Linux workstations equipped with Nvidia Quadro RTX 5000 graphics processing units (GPUs) and a compute cluster with Nvidia V100 and A100 GPUs. In case of the DNNs, we used the Adam optimizer to accelerate learning. Parameter values used in all simulations are summarized in Supplementary Table S3.

### Learning in the single neuron setup

We considered a simple linear rate-based neuron model whose output firing rate *z* is given by the weighted sum of the firing rates *x_j_* of the input neurons, i.e, *z* = ∑_*j*_ *W_j_x_j_*, where *W_j_* corresponds to the synaptic weight of input *j*. We trained the neuron using stochastic gradient descent (SGD) on the corresponding objective function:

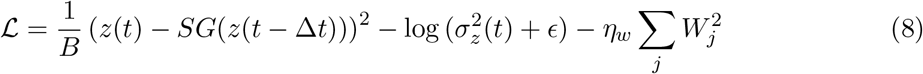

Here, and in all following simulations, we fixed the Hebbian coefficient λ_1_ = 1. We also added a small constant *ϵ* = 10^-6^ to the estimate of the variance *σ_z_* for numerical stability. In case of a single rate neuron, the LPL rule (Eq. (7)) simplifies to Eq. (1) without the decorrelation term.

#### Synthetic two-dimensional dataset generation

The two-dimensional synthetic data sequence (Fig. 2a) consists of two clusters of inputs, one centered at *x* = −1, and the other at *x* = +1. Pairs of consecutive data points were drawn independently from normal distributions centered at their corresponding cluster. To generate a family of different datasets, we kept the standard deviation in the x-direction fixed at *σ_x_* = 0.1 and varied *σ_y_*. Additionally, to account for occasional transitions between clusters with probability *p*, we included a corresponding fraction of such “crossover pairs” in the training batch. For each value of *σ_y_*, we simulated the evolution of the input connections of a single linear model neuron that received the *x* and *y* as its two inputs, and updated its input weights according to LPL. In the simulations in Fig. 2 we assumed *p* → 0, however, the qualitative behavior remained unchanged for noise levels below *p* = 0.5, i.e, as long as the “noisy” pairs of points from different clusters were rare in each training batch (Extended Data Fig. 10).

#### Neuronal selectivity measure

After training weights to convergence, we measured the neuron’s selectivity to the x-input as the normalized difference between mean responses to stimuli coming from the two respective input clusters. Concretely, let 〈*z*_1_〉 be the average output caused by inputs from the *x* = 1 cluster, and 〈*z*_2_〉 from the *x* = –1 cluster, then the selectivity *χ* is defined as:

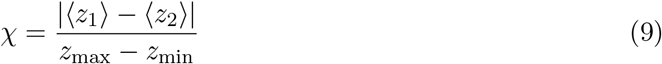

with *z*_max_ the maximum and z_min_ the minimum response across all inputs.

### Learning in deep convolutional neural networks

For all network simulations, we used a convolutional DNN based on the VGG-11 architecture [55] (see Supplementary Note S5 for details). We trained this network on STL-10 and CIFAR-10 (Extended Data Fig. 11), two natural image datasets (see Supplementary Table S3 for hyperparameters). To simulate related consecutive inputs, we used two differently augmented versions of the same underlying image, a typical approach in vision-based SSL methods. Specifically, we first standardized the pixel values to zero mean and unit standard deviation within each dataset before using the set of augmentations originally suggested in [21], which includes random crops, blurring, color jitter and random horizontal flips (see Extended Data Fig. 2 for examples).

#### Synthetic video generation

To study LPL in settings with more naturalistic transitions between consecutive images and without relying on image augmentation, we procedurally generated videos using images from the 3D Shapes dataset [42]. The dataset has a known latent manifold structure spanned by view angle, object scale, hue, and object type and is commonly used to measure disentangling in variational autoencoders. Using the knowledge of the ground truth factors, we generated continuous video composed of 17-frame clips during which the object shape remained fixed and a randomly chosen factor changed gradually. Specifically, we proceeded as follows: we randomly chose one factor and changed it frame-by-frame such that transitions between adjacent factor values were more likely. For instance, one such clip shows a cube under a smoothly varying camera angle (Extended Data Fig. 5a). Furthermore, we randomly permuted the order of all three hue factors. This was done to break the orderly ring topology of the hue mappings in the original dataset, which allowed us to test that the structure is restored through LPL, but not other methods (see Extended Data Fig. 5g). After 17 frames we randomly chose another shape and factor and repeated the above procedure. This sequence generation resulted in video with many consecutive latent manifold traversals as captured by the empirical transition matrices (Extended Data Fig. 6a). Importantly, due to the nature of the video, which switches between objects periodically, the resulting input sequence also included occasional transitions between different objects that the LPL rule interprets as positive samples. Such transitions also appear in real world stimuli when objects leave or enter the scene. Despite these “false positives” LPL learned disentangled representations of shapes and the underlying factors.

#### Network training

We trained our network models on natural image data by minimizing the equivalent LPL objective function. For both datasets, we trained the DNN using the Adam optimizer with default parameters and a cosine learning rate schedule that drove the learning rate to zero after 800 epochs. We distinguish between two cases: layer-local and end-to-end learning. End-to-end learning corresponds to training the network by optimizing 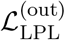 at the network’s output while using backpropagation to train the hidden layer weights. This is the standard approach used in deep learning. In contrast, in layer-local learning, we minimized the LPL objective 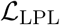 at each layer in the network independently without backpropagating loss gradients between layers similar to previous work [24, 31]. In this case, every layer greedily learns predictive features of its own inputs, i.e, its previous layer’s representations. To achieve this behavior, we prevented PyTorch from backpropagating gradients between layers by detaching the output of every layer in the forward pass and optimizing the sum of per-layer losses 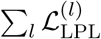.

Unless mentioned otherwise, we used global average pooling (GAP) to reduce feature maps to a single vector before applying the learning objective at the output of every convolutional layer for layer-local training, or just at the final output in the case of end-to-end training. Although pooling was not strictly necessary and LPL could be directly applied on the feature maps, it substantially sped up learning and led to an overall improved linear readout accuracy on CIFAR-10 (Supplementary Table S2). However, we observed that GAP was essential on the STL-10 dataset for achieving readout accuracy levels above the pixel-level baseline (cf. Table 1). This discrepancy was presumably due to the larger pixel dimensions of this dataset and the resulting smaller relative receptive field size in early convolutional layers. Concretely, feature pixels in the first convolutional layer of VGG-11 have a receptive field of 3 × 3 pixels covering a larger portion of the 32 × 32 CIFAR-10 images as compared to the 96 × 96 STL-10 inputs. This hypothesis was corroborated by the fact that when we sub-sampled STL-10 images to a 32 × 32 resolution, the dependence on GAP was removed and LPL was effective directly on the feature maps (Supplementary Table S2).

#### Baseline models

As baseline models for comparison (Supplementary Table S1), we trained the same convolutional neural network (CNN) network architecture either with a standard crossentropy supervised objective, which requires labels, or with a contrastive objective, which relies on negative samples. To implement contrastive learning, the network outputs *z*(*t*) were passed through two additional dense projection layers *v*(*t*) = *f*_proj_(*z*(*t*)), which is considered crucial in contrastive learning to avoid dimensional collapse [41]. Finally, the following contrastive loss function was applied to these projected outputs

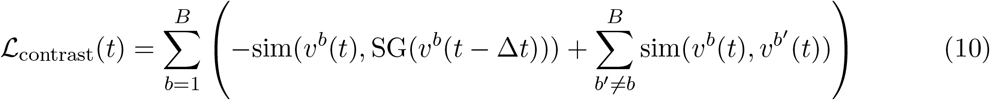

where 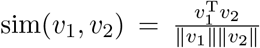 is the cosine similarity between two representations *v*_2_ and *v*_2_. The second term in the loss function is a sum over all pairwise similarities between inputs in a given minibatch. These pairs correspond to different underlying base images and therefore constitute negative samples. During training the network is therefore optimized to reduce the representational similarity between them.

For training the layer-local versions of the supervised and contrastive models, we followed the same procedure as with LPL of optimizing the respective loss function at the output of every convolutional layer *l* of the DNN without backpropagation between the layers. Because projection networks are necessary for avoiding dimensional collapse in case of contrastive learning, we included two additional dense layers to obtain the projected representations 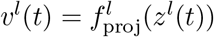 at every level of the DNN before calculating the layer-wise contrastive loss 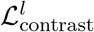. This meant that gradients were backpropagated through each of these dense layers for training the corresponding convolutional layers of the DNN, but consecutive convolutional layers were still trained independent of each other.

#### Population activity analysis

We adopted two different metrics in order to analyze the representations learned by the DNN after unsupervised training with LPL on the natural image datasets.

#### Linear readout accuracy

To evaluate how well the LPL rule trained the DNN to disentangle and identify underlying latent factors in a given image, we measured linear decodability by training a linear classifier on the network outputs in response to a set of training images. Crucially, during this step we only trained the readout weights while keeping the weights of the LPL-pretrained DNN frozen. We then evaluated the linear readout accuracy (Fig. 3b) on a held-out test set of images. We used the same procedure to evaluate the representations at intermediate layers (Fig. 3c), and for the baseline models.

#### Representational similarity analysis

To visualize the latent manifold structure in learned network embeddings, we computed average representational similarity matrixs (RSMs). To obtain the RSM for one factor, say object hue, we first fixed the values of all the other factors and calculated the cosine similarity between the network outputs as the object hue was changed. We repeated this procedure for many different values for the other factors to get the final averaged RSM for object hue (Extended Data Fig. 5f).

#### Metric for disentanglement

To quantitatively measure disentanglement, we used the metric proposed by Kim and Mnih [42]. This measure requires full knowledge of the underlying latent factors, as was the case for our procedurally generated videos. In brief, to compute the measure one first identifies the most insensitive neuron to all except one factor. Next, using the indices of these neurons, one trains a simple majority-vote classifier that predicts which factor is being coded for. The accuracy of this classifier on held-out data is the disentanglement score.

#### Dimensionality and activity measures

To characterize mean activity levels in the network models, we averaged neuronal responses over all inputs in the validation set. To quantify the dimensionality of the learned representations, we computed the participation ratio [56]. Concretely, if 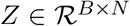 are *N*-dimensional representations of *B* input images, and λ_*i*_, 1 ≤ *i* ≤ *N* is the set of eigenvalues of *Z^T^Z*, then the participation ratio is given by:

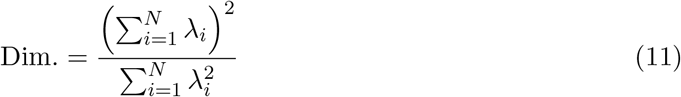

### Model of unsupervised learning in inferotemporal cortex

#### Network model and pretraining dataset

To simulate the experimental setup of Li and DiCarlo [14], we modeled the animal’s ventral visual pathway with a convolutional DNN. To that end, we used the same network architecture as before except that we removed all biases in the convolutional layers in order to prevent boundary effects. This modification resulted in a drop in linear readout accuracy (Supplementary Table S2). Pre-training of the network model proceeded in two steps as follows. First, we performed unsupervised pre-training for 800 epochs on STL-10 using augmented image views exactly as before. Next, we added a fully-connected dense layer at the network’s output, and trained it for 10 epochs with the LPL objective while keeping the weights of the convolutional layers frozen. During this second pre-training phase, we used augmented STL-10 inputs which were spatially extended in order to account for the added spatial dimension of different canvas positions in the experiment [14]. The expanded inputs consisted of images placed on a large black canvas at either the center position *X_c_* or one of two peripheral positions *X*_1/2_ at the upper or lower end of the canvas. Concretely, these images had dimensions (13 × 96) × 96 which resulted in an expanded feature map at the output of the convolutional DNN with spatial dimensions 13 × 1 (see Supplementary Note S5 for details). Note that we only expanded the canvas in the vertical dimension instead of using a setup with a 13 × 13 feature map because it resulted in a substantial reduction of computational and memory complexity. During this second stage of pre-training, the network was only exposed to “true” temporal transitions wherein the image was not altered between time steps apart from changing position on the canvas.

#### Data generation for simulated swap exposures

To simulate the experiment by [14], we exposed the network to normal and swap temporal transitions. In the latter case the image was consistently switched to one belonging to a different object category at the specific swap position. The swap position for a given pair of images was randomly pre-selected to be either *X*_1_ or *X*_2_, while the other non-swap position was used as a control. Specifically, we switched object identities during transitions from one peripheral swap position, say *X*_1_, to the central position *X_c_*, while keeping transitions from the other peripheral position *X*_2_ to the center unmodified. As in the experiment, we chose several pairs of images as swap pairs, and fixed *X*_1_ as the swap position for half the pairs of images and *X*_2_ as the swap position for the other half. To simulate ongoing learning during exposure to these swap and non-swap input sequences, we continued fine-tuning the convolutional layers. To that end, we used the Adam optimizer we used during pre-training with its internal state restored to the state at the end of pre-training. Moreover, we used a learning rate of 10^-7^ during fine-tuning which was approximately 100 × larger than the learning rate reached by the cosine learning rate schedule during pre-training (4 × 10^-9^, after 800 epochs). Finally, we trained the newly added dense layers with vanilla SGD with a learning rate of 0.02.

#### Neuronal selectivity analysis

Before training on the swap exposures, for each output neuron in the dense layer, we identified the preferred and non-preferred member of each swap image pair, based on which image drove higher activity in that neuron. This allowed us to quantify object selectivity on a per-neuron basis as *P* – *N*, where *P* is the neuron’s response to its initially preferred image, and *N* to its nonpreferred image at the same position on the canvas. Note that, by definition, the initial object selectivity for every neuron is positive. Finally, we measured the changes in object selectivity *P* – *N* during the swap training regimen, at the swap and non-swap positions averaging over all output neurons for all image pairs. As a control, we included measurements of the selectivity between pairs of control images that were not part of the swap set.

#### Comparison to experimental data

To compare our model to experiments, we extracted the data from [14] using the Engauge Digitizer software and replotted it in Fig. 4b.

### Spiking neural network simulations

We tested a spiking version of LPL in networks of conductance-based leaky integrate-and-fire (LIF) neurons. Specifically, we simulated a recurrent network of 125 spiking neurons (100 excitatory and 25 inhibitory neurons) receiving afferent connections from 500 input neurons. In all simulations the input connections evolved according to the spike-based LPL rule described below. In our model, neurons actively decorrelated each other through locally connected inhibitory interneurons whose connectivity was shaped by inhibitory plasticity.

#### Neuron model

The neuron model was based on previous work [28, 57] in which the membrane potential *U_i_* of neuron *i* evolves according to the ordinary differential equation

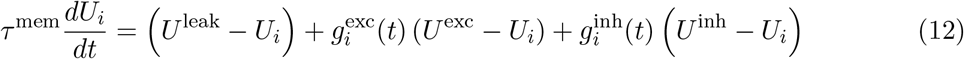

where *τ*^mem^ denotes the membrane time constant, *U^x^* are the synaptic reversal potentials (Supplementary Table S4), and the 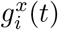 the corresponding synaptic conductances expressed in units of the neuronal leak conductance. The excitatory conductance is the sum of NMDA and AMPA conductances 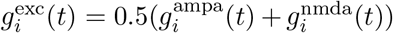. Their dynamics are described by the following differential equations

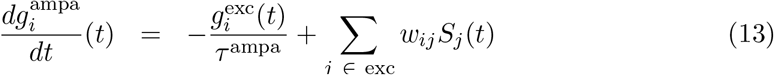

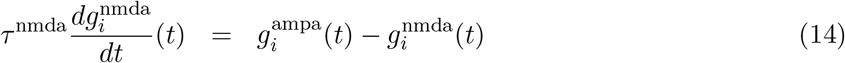

whereas the inhibitory GABA conductance 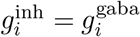 evolves as

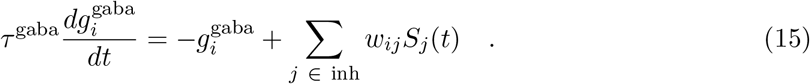

In the above expressions 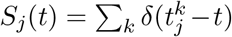 refers to the afferent spike train emitted by neuron *j*, in which 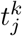 are the corresponding firing times, and *τ^x^* denotes the individual neuronal and synaptic time constants (Supplementary Table S4). Neuron *i* fires an output spike whenever its membrane potential reaches the dynamic firing threshold *ϑ_i_*(*t*) that evolves according to

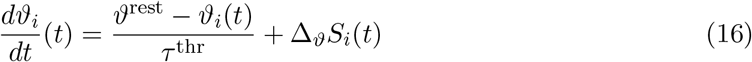

to implement an absolute and relative refractory period. Specifically, *ϑ_i_* jumps by Δ_*ϑ*_ = 100 mV every time an output spike is triggered after which it exponentially decays back to its rest value of *ϑ*^rest^ = –50 mV. All neuronal spikes are delayed by 0.8 ms to simulate axonal delay and to allow efficient parallel simulation before they trigger postsynaptic potential in other neurons.

#### Time varying spiking input model

Inputs were generated from 500 input neurons divided into five populations of 100 Poisson neurons each. All inputs where implemented as independent Poisson processes with the same average firing rate of 5 Hz and neurons within the same group shared the same instantaneous firing rate. Concretely, neurons in *P*0 had a fixed firing rate of 5 Hz, whereas the firing rates in groups *P*1 and *P*2 changed slowly over time. Specifically, we generated periodic template signals *x*(*t*) from a Fourier basis

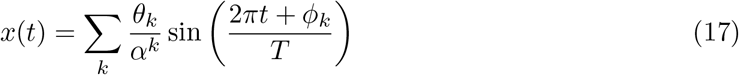

with random uniformly drawn coefficients 0 ≤ *θ_k_, ϕ_k_* < 1. The spectral decay constant *α* = 1.1 biased the signals toward slow frequencies and thus slowly varying temporal structure. We chose the period *T* = 3 s for *P*1 and (3+^1/13^)s for *P*2 respectively. The different periods were chosen to avoid phase-locking between the two signals. Both signals were then sampled at 10 ms intervals, centered on 5 Hz, variance-normalized, and clipped below at 0.1Hz before using them as periodic time varying firing rates for *P*1 and *P*2. Additionally, we simulated control inputs *P*1/2_ctl_ of the two input signals by destroying their slowly varying temporal structure. To that end, we repeated the original firing rate profile for 13 periods before shuffling it on a time grid with 10 ms temporal resolution.

#### Spike-based LPL

To extend LPL to the spiking domain, we build on SuperSpike [58], a previously published online learning rule, which had only been used in the context of supervised learning in SNNs thus far. In this article, we replaced the supervised loss with the LPL loss (Eq. (3)) without the decorrelation term. The resulting spiking LPL online rule for the weight *w_ij_* is given by

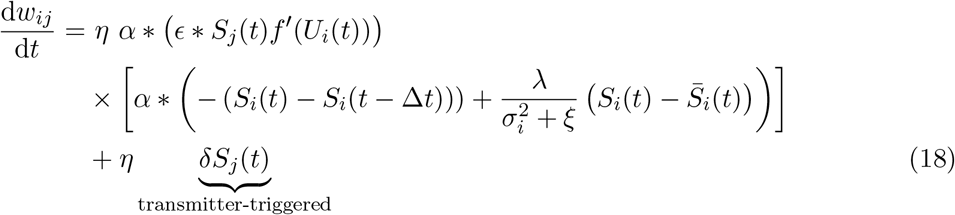

with the learning rate *η* = 10^-2^, a small positive constant *ξ* = 10^-3^ to avoid division by zero. Further, *α* is a double exponential causal filter kernel applied to the neuronal spike train *S_i_*(*t*). Similarly, *ϵ* is a causal filter kernel that captures the temporal shape of how a presynaptic spike influences the postsynaptic membrane potential. For simplicity, we assumed a fixed kernel and ignored any conductance-based effects and NMDA dependence. Further, we added the transmitter-triggered plasticity term with *δ* = 10^-5^ to ensure that weights of quiescent neurons slowly potentiate in the absence of activity to ultimately render them active [57]. Finally, λ = 1 is a constant that modulates the strength of the Hebbian term. We set it to zero to switch off the predictive term where this is mentioned explicitly.

Further, *f*′(*U_i_*) = *β* (1 + *β*|*U_i_* – *ϑ*^rest^|) is the surrogate derivative with *β* = 1 mV^-1^, which renders the learning rule voltage-dependent. Finally, 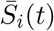 and 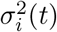 are slowly varying quantities obtained online as exponential moving averages with the following dynamics

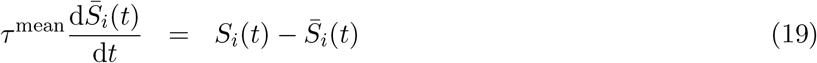

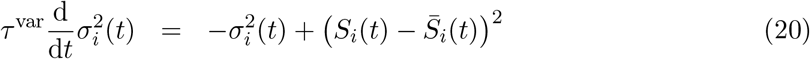

with *τ*^mean^ = 600 s and *τ*^var^ = 20 s. These quantities confer the spiking LPL rule with elements of metaplasticity [34].

In our simulations, we computed the convolutions with *α* and *ϵ* by double exponential filtering of all quantities. Generally, for the time varying quantity *c*(*t*) we computed

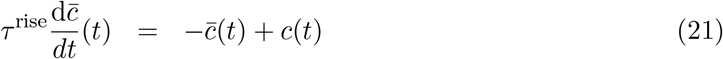

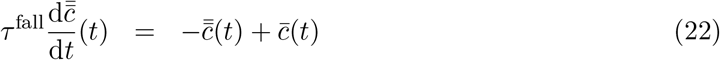

which yields the convolved quantity 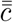. Specifically, we used 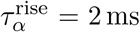, 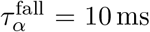, 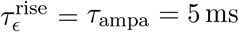, and 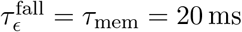.

Overall, one can appreciate the resemblance of Eq. (18) with the non-spiking equivalent (cf. Eq. (1)). As in the non-spiking case the learning rule is local in that it only depends on pre- and postsynaptic quantities. The predictive term in the learning rule can be seen as an instantaneous error signal which is minimized when the present output spike train *S_i_*(*t*) is identical to a delayed version of the same spike train *S_i_*(*t* – Δ*t*) with Δ*t* = 20 ms. In other words, the past output serves as a target spike train (cf. 58).

#### Microcircuit connectivity

Connections from the input population to the network neurons and recurrent connections were initialized with unstructured random sparse connectivity with different initial weight values (Supplementary Table S5). One exception to this rule was the excitatory-to-inhibitory connectivity which was set up with a Gaussian connection probability profile

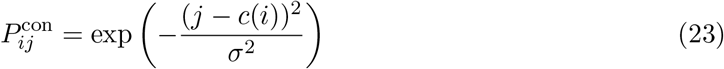

with *c*(*i*) = 0.25*i* with *σ*^2^ = 20 to mimic the dense local connectivity onto inhibitory neurons due to which inhibitory neurons inherit some of the tuning of their surrounding excitatory cells.

#### Inhibitory plasticity

Inhibitory to excitatory synapses were plastic unless mentioned otherwise. We modeled inhibitory plasticity according to a previously published inhibitory STDP model [38].

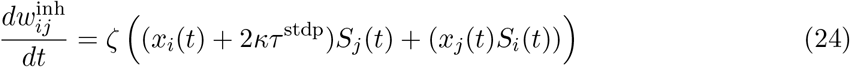

using pre- and postsynaptic traces

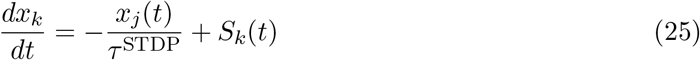

with time constant *τ*^STDP^ = 20 ms, learning rate *ζ* = 1 × 10^-3^, and target firing rate *κ* = 10 Hz.

#### Reconstruction of input signals from network activity

To reconstruct the input signals, we first computed input firing rates of the five input populations by binning their spikes emitted during the last 100 s of the simulation in 25 ms bins. We further averaged the binned spikes over input neurons to provide the regression targets. Similarly, we computed the binned firing rates of the network neurons but without averaging over neurons. We then performed Lasso regression using SciKit-learn with default parameters to predict each target input signal from the network firing rates. Specifically, we trained on the first 95 s of the activity data, and computed *R*^2^ scores on the Lasso predictions over the last 5 s of held-out data (Fig. 5b).

#### Signal selectivity measures

We measured signal selectivity of each neuron to the two slow signals relative to their associated shuffled controls (Fig. 5d) using the following relative measure defined on the weights:

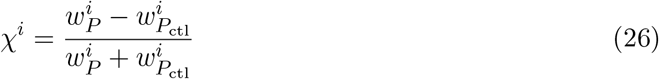

where 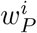 is the average synaptic connection strength from the signal pathways *P*1/2 onto excitatory neuron *i*, and 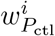 is the same but from the control pathways *P*1/2_ctl_.

#### Representational dimension

To quantify the dimensionality of the learned neuronal representations (Fig. 5f), we binned network spikes in 25 ms bins and computed the participation ratio (Eq. (11)) of the binned data.

#### Neuronal tuning analysis of the learned weight profiles

To characterise the receptive fields of each neuron (Fig. 5g,h), we plotted *w*_*P*1_ against *w*_*P*2_ for every neuron in the excitatory population (Figs. 5g,h; left), and colored the resulting weight vectors by mapping the cosine of the vectors with the x-axis (*w*_*P*2_) to a diverging color map. Furthermore, we calculated the relative tuning index as follows

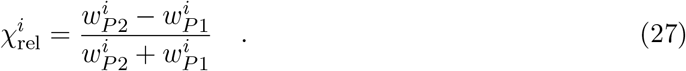

#### STDP induction protocols

To measure STDP curves, we simulated a single neuron using the spiking LPL rule (Eq. 18) with a learning rate of *η* = 5 × 10^-3^. In all cases, we measured plasticity outcomes from 100 pairings of pre- and postsynaptic spikes at varying repetition frequencies *ρ*. The postsynaptic neuron’s membrane voltage was held fixed between spikes at −51mV for the entire duration of the protocol. To measure STDP curves, we set the initial synaptic weight at 0.5 and simulated 100 different pre-post time delays Δ*t* chosen uniformly from the interval [−50, 50] ms with *ρ* = 10 Hz. To measure the rate-dependence of plasticity, we repeated the simulations for fixed Δ*t* = ±10 ms while varying the repetition frequency *ρ*.

#### Numerical simulations

All SNN simulations were implemented in C++ using the Auryn SNN simulator. Throughout we used a 0.1 ms simulation time step. Simulations were run on seven Dell Precision workstations with eight-core Intel Xeon CPUs.

## Data availability

The deep learning tasks used the STL-10 and CIFAR-10 datasets, typically available through all major machine learning libraries. The original releases for these datasets can be found at http://ai.stanford.edu/%7Eacoates/stl10/, and https://www.cs.toronto.edu/~kriz/cifar.html respectively. We further used the 3D Shapes dataset [42] available at https://github.com/deepmind/3d-shapes/.

## Code availability

- The simulation code to reproduce the key results is publicly available at https://github.com/fmi-basel/latent-predictive-learning.
- PyTorch and the Lightning framework are freely available at https://pytorch.org and https://www.pytorchlightning.ai.
- The Auryn spiking network simulator is available at https://github.com/fzenke/auryn.
- The Engauge Digitizer is available at http://markummitchell.github.io/engauge-digitizer.

## Acknowledgments

We thank all members of the Zenke Group for comments and discussions that shaped this project, and Atul Kumar Sinha for many helpful suggestions. We are particularly grateful to Julian Rossbroich for providing invaluable insights throughout the course of this work. This project was supported by the Swiss National Science Foundation [grant number PCEFP3_202981] and the Novartis Research Foundation.

## Author contributions

F.Z. conceived the study. M.S.H. and F.Z. developed the theory. M.S.H. wrote DNN code, performed simulations, and analysis. F.Z. developed SNN code. M.S.H. and F.Z. wrote the manuscript.

---

**Extended Data Fig. 1:**
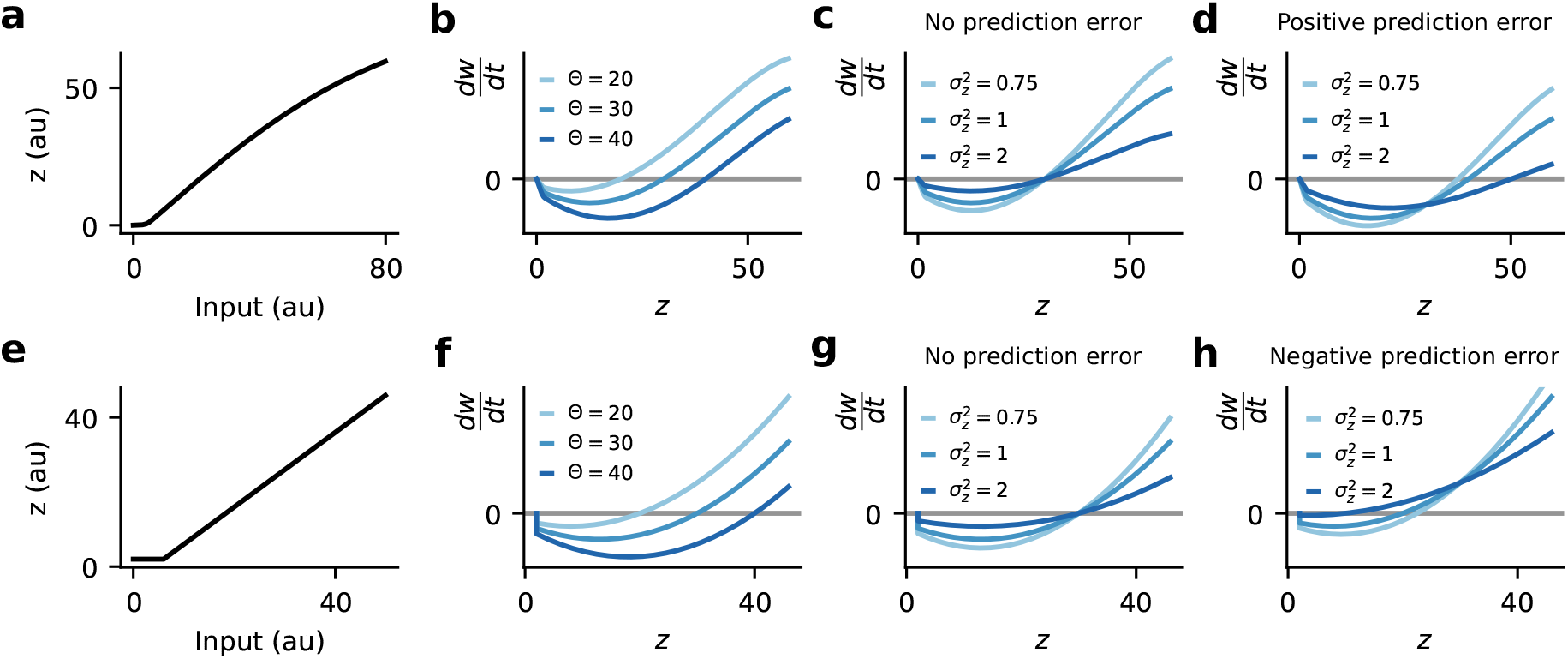
LPL extends BCM theory by adding a variance- and rate-of-change dependence. **(a)** Example of a typical neuronal input-output function with postsynaptic activity *z*. **(b)** Weight change induced by the LPL rule for co-varying input and the postsynaptic activity *z* for different values of the plasticity threshold Θ, with 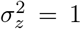 and 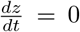. The functional shift of the threshold is reminiscent of the BCM rule. **(c)** Same as (b) but for different values of the variance of the postsynaptic activity with zero prediction error 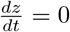 and fixed mean activity 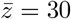. **(d)** Same as (c) but with a positive prediction error 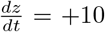. **(e)** Same as (a), but for a rectified linear unit (ReLU) activation function with positive threshold. **(f–g)** Same as above but for ReLU. **(h)** Same as in (d) but for ReLU and a negative prediction error 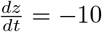.

**Extended Data Fig. 2:**
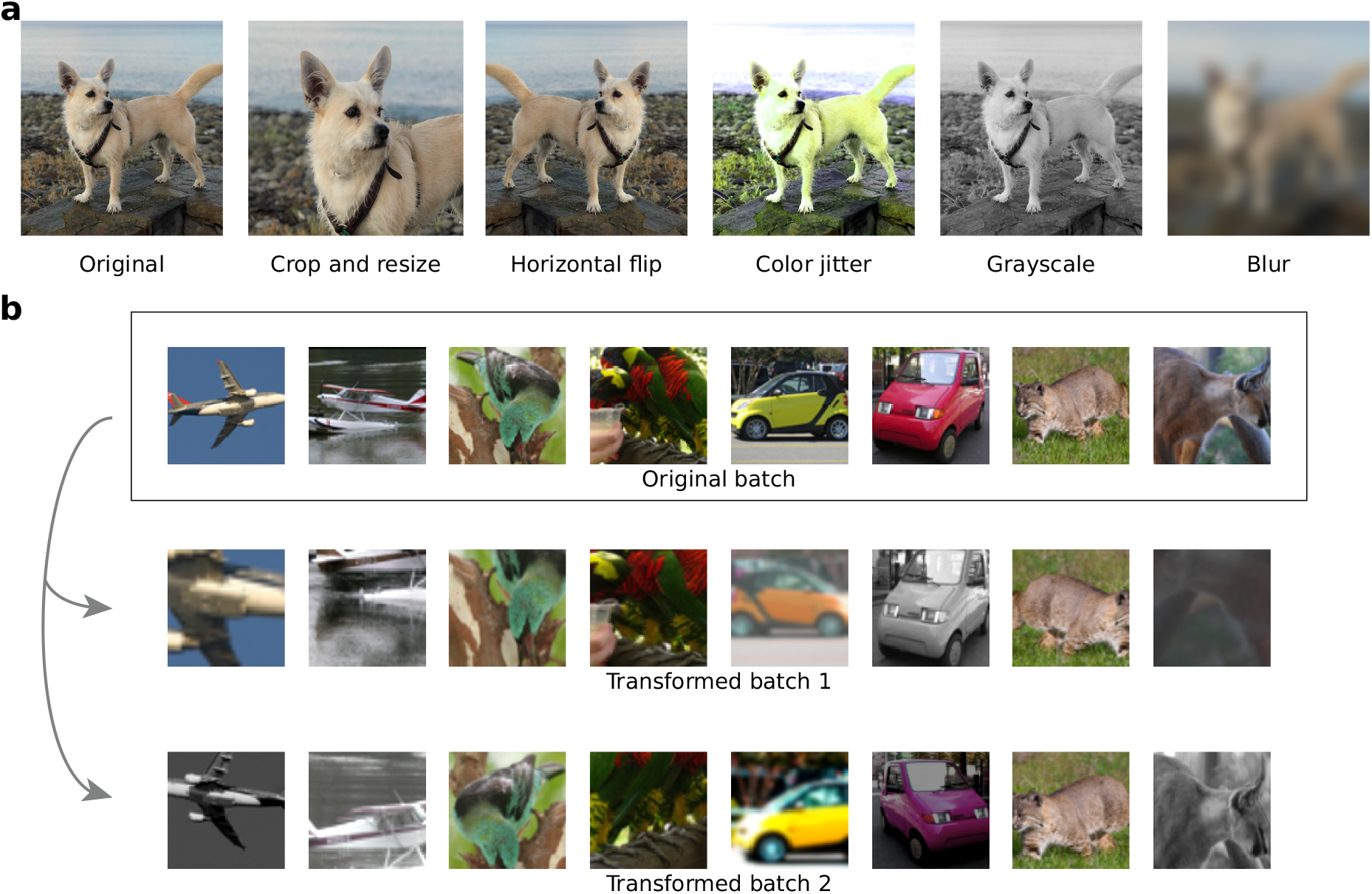
Image augmentation model. **(a)** Illustration of the image transformations used to generate natural image sequences as suggested by Chen *et al* [21]. **(b)** Sample images from STL-10 and their transformed versions used for training the DNNs.

**Extended Data Fig. 3:**
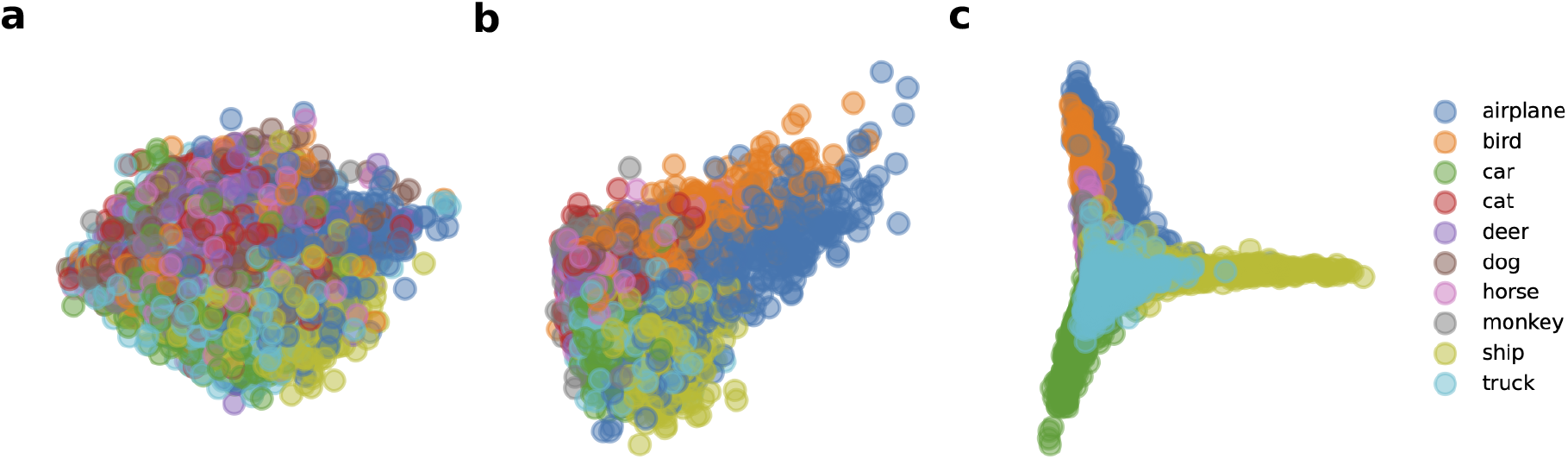
Disentangling of object representations in the DNN. **(a)** Data distribution of the STL-10 validation set along the first two principal components in pixel-space. Data corresponding to different object classes are highly entangled. **(b)** Same as (a) but along the principal components of representations in Layer 3 of the DNN after learning with LPL. Object classes are somewhat disentangled. **(c)** Same as (a) but along the principal components of representations in Layer 8 of the DNN. Object classes are highly disentangled.

**Extended Data Fig. 4:**
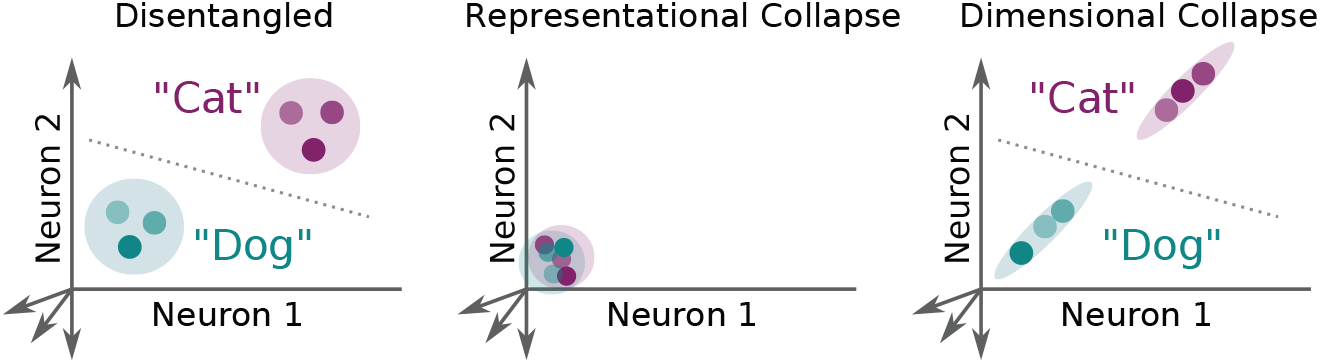
Illustration of collapse modes typifying poorly disentangled features. Effectively disentangled representations (left) separate categories well with different representational directions encoding different relevant features. Purely predictive learning without counteracting Hebbian plasticity leads to collapsed representations (center), typically to zero activity levels. Dimensional collapse (right) is characterized by highly correlated activity across all neurons, indicating that only one relevant feature is encoded by all neurons, which is unlikely to be conducive to hierarchical feature extraction for non-trivial tasks such as object recognition.

**Extended Data Fig. 5:**
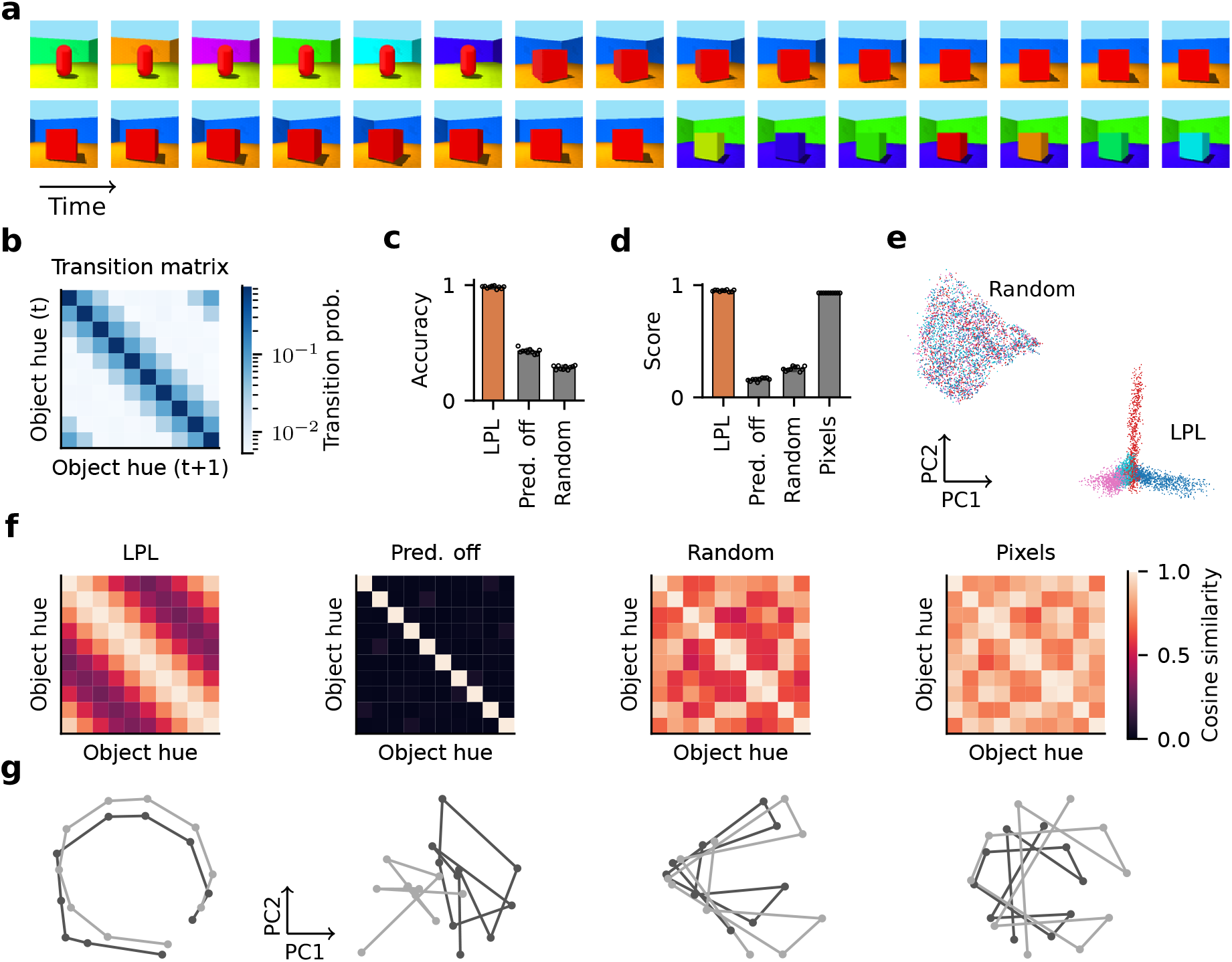
LPL finds latent manifold structure of simulated video data. **(a)** Input frames from the procedurally generated video using the 3D Shapes dataset [42]. **(b)** The empirically measured transition matrix of object hue with latent structure (see Extended Data Fig. 6 for the complete set of transition matrices). **(c)** Object classification accuracy of a linear classifier trained on network outputs of a network with LPL, without the predictive term (Pred. off), and the randomly initialized network (Random). Values represent averages from ten-fold cross validation. The accuracy is close to 100% for LPL, but lower at initialization or when trained without the predictive term. **(d)** Disentanglement scores computed according to the metric proposed by Kim and Mnih [42] for the final-layer representations of the three networks in (c) compared to the input pixels (Pixels). LPL yields close to maximum scores (95.0% ± 0.8%), higher than a randomly initialized network or after training without the predictive term. However, evaluating the metric on the pixels directly also yields high scores (93.0% ± 0.0%), albeit still slightly lower than LPL. The high scores in pixel space can partially be explained by the high input dimension and the small number of classes in the dataset. Importantly, the metric is insensitive to the manifold topology (see below). Different data points correspond to averaging over ten independent evaluations of the metric. **(e)** Projections of the representations onto the first two principal components before (Random) and after training (LPL). Each point corresponds to one input image, and the color represents the object type. The object class is entangled at initialization and disentangled after learning. **(f)** Averaged RSM computed from representations of different object colors in (d). LPL’s RSM closely resembles the transition structure shown in (b). Without the predictive term, the RSM becomes diagonal, while the random network’s RSM does not have this structure and roughly follows the input pixel similarity structure. **(g)** Network output projected onto the first two principal components for changing hue sequentially while keeping all other factors fixed. The two lines correspond to two different object sizes. The trajectories are disentangled for LPL and preserve the topology of the data manifold (cf. b), whereas this is not the case when the predictive term is off, at initialization (random), or at the input (pixels).

**Extended Data Fig. 6:**
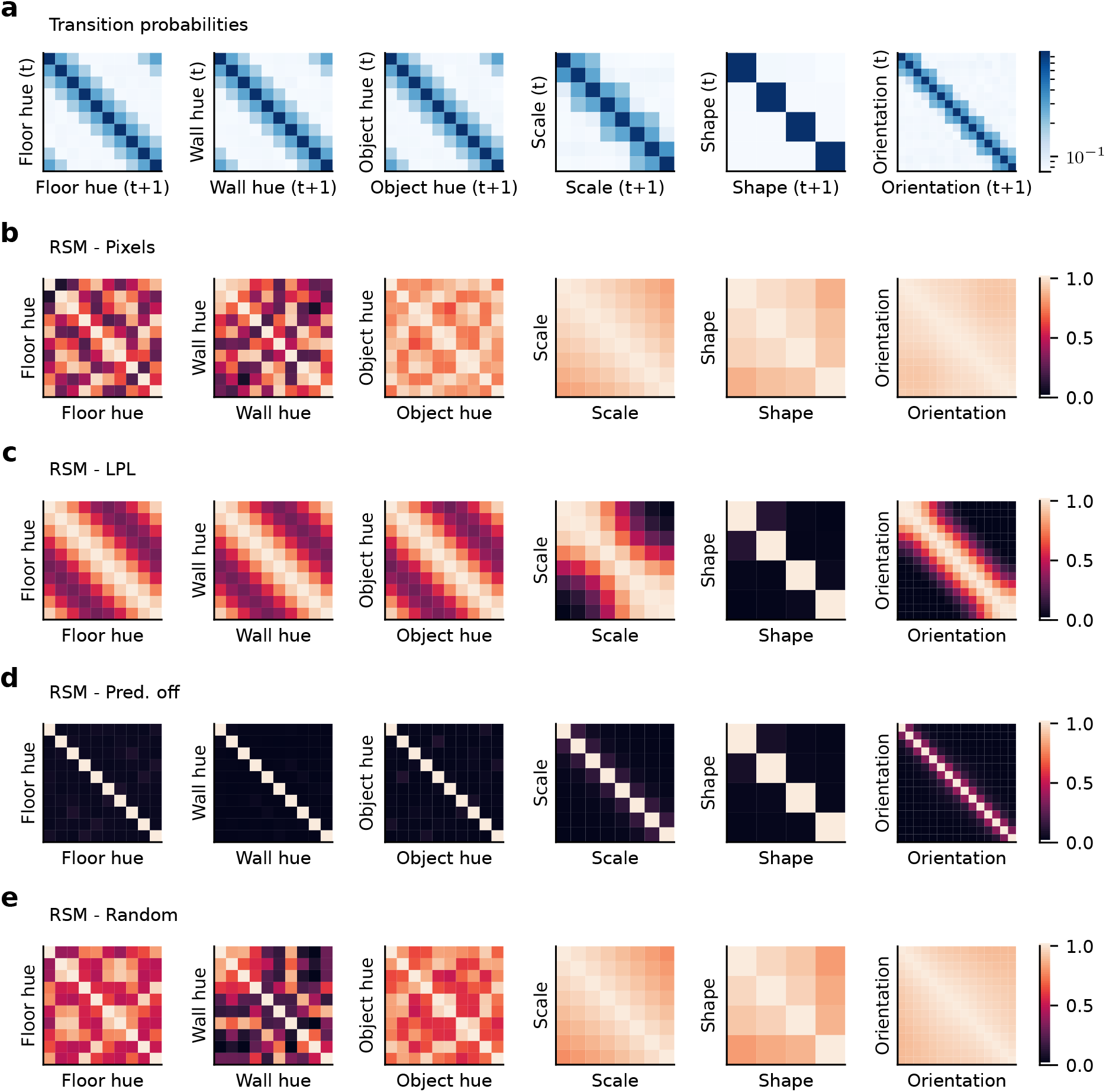
Transition matrices and RSMs for all latent factors of the 3D shapes dataset. **(a)** Transition probabilities estimated from the generated video. The high values on the diagonal reflect the fact that within a 17-frame clip, only one factor changes while the others remain fixed. The off-diagonal values reflect the transition probabilities when a specific factor is changing. For instance, within a clip cycling through all the object hues, the color may only change to the next or previous assignments in the color map with a smaller probability for a two-step transition. The hue mapping was randomly chosen with respect to the original dataset to ensure an entangled topology at the input (cf. Fig. 5g). The orientation and scale factors are not allowed to transition from the smallest to the largest values, and vice versa. Furthermore, the direction of change for any factor is fixed within a given clip, but may reverse for orientation and scale at the extreme values (cf. Fig. 5a). **(b)** Same as Fig. 5f, but for all factors at the pixel level. RSM values represent average cosine similarity between the pixels of images differing only in one factor with all other factors fixed. Some similarity structure exists along the scale and orientation factors only. **(c)** Same as (b) but for the final-layer representations learned by LPL. The RSM closely resembles the transition probability structure that characterizes the temporal properties of the video sequence. **(d)** Same as (c) but for learning without the predictive term. The RSM is diagonal, which shows that the network represents different factors in almost orthogonal directions. **(e)** Same as (b), but at random initialization before training. The RSM for all factors is reflective of the pixel RSM.

**Extended Data Fig. 7:**
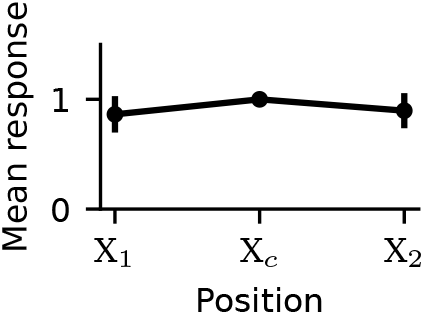
Learned network representations are invariant to object position on the canvas. Activity of neurons in the pretrained DNN’s output layer in response to images at three positions on the canvas, normalized by each neuron’s response to the center position, and averaged over neurons and over images.

**Extended Data Fig. 8:**
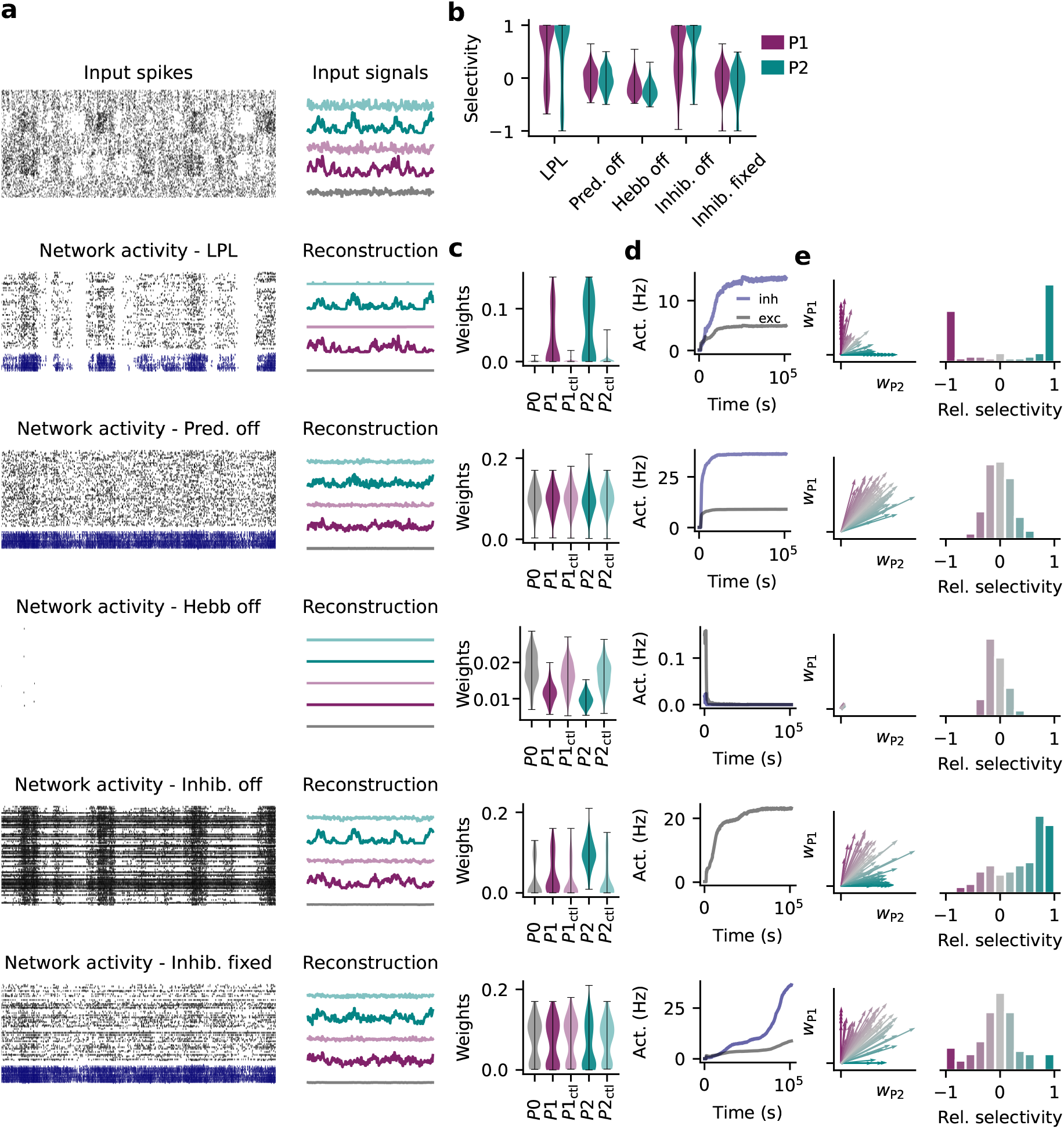
Same as Figure 5 but with detailed controls. **(a)** Snapshot of spiking activity (left) and underlying firing rate signals or their reconstructions (right) over 100 ms for the input and network populations. **(b)** Same as Fig. 5d showing signal selectivity learned with the different variations of spiking LPL given in (a). **(c)** Average synaptic connection strength grouped by input population for the different configurations in (a). LPL with plastic inhibition results in higher weights on the slowly varying signals relative to the shuffled controls, but not when the predictive or Hebbian term are disabled. Without inhibition or without inhibitory plasticity, connections from all populations are strong with a small preference for *P*2. **(d)** Average firing rates over 100 s bins throughout training for the configurations in (a). Firing rates saturate with the inhibitory neurons settling at a higher firing rate when learning with spiking LPL with inhibition, even when the predictive term is disabled or the inhibition is not plastic. Activity collapses without the Hebbian term, whereas firing rates diverge without inhibition. **(e)** Averaged weight vectors from populations *P*1 and *P*2 onto each excitatory neuron (left) and distribution of the excitatory neurons’ relative selectivity between the two populations (right). Different neurons are exclusively selective to either *P*1 pr *P*2 under spiking LPL with inhibitory plasticity. Without the predictive term, or the Hebbian term, few if any neurons are selective to one population over the other. Moreover, weights collapse to small values without the Hebbian term. When inhibition is removed altogether, a few neurons become exclusively selective to *P*2, but the weight vectors are not well-decorrelated. Without inhibitory plasticity, a few weight vectors are well-decorrelated, but most neurons are not preferentially selective to either signal.

**Extended Data Fig. 9:**
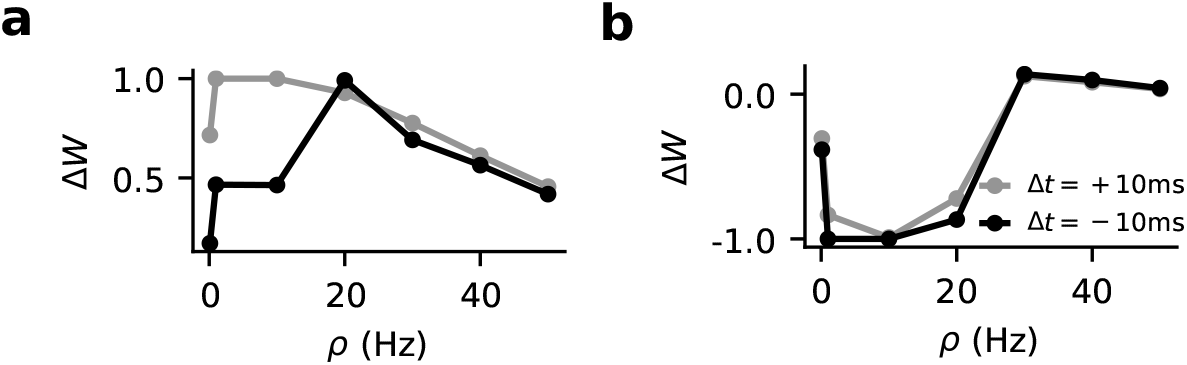
Learning threshold determines the sign of plasticity. **(a)** Weight changes as a function of repetition frequency *ρ* for positive and negative relative spike timings (Δ*t* = ±10 ms) with *σ*^2^(*t* = 0) = 0 and 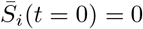. **(b)** Same as (a) but for 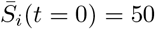.

**Extended Data Fig. 10:**
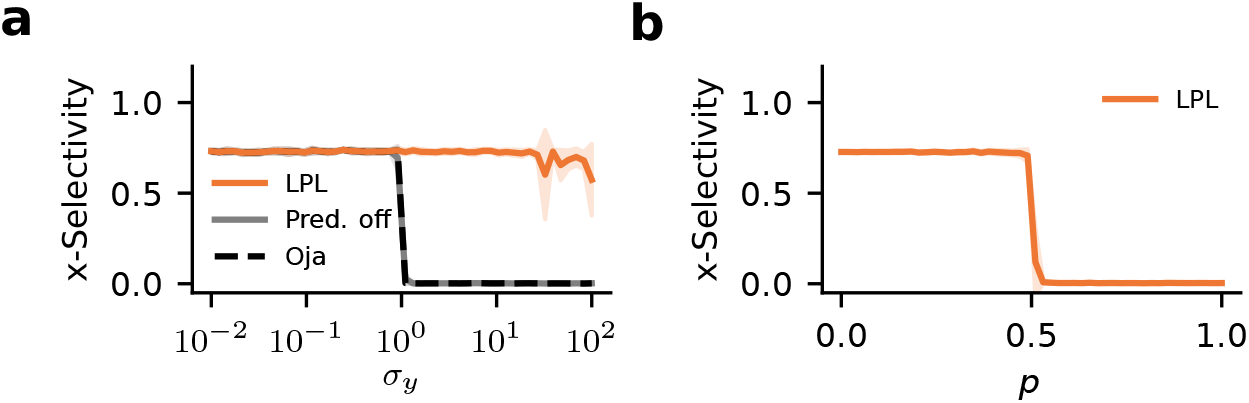
LPL is robust to noise. **(a)** Same as Figure 1b but for high rates of noisy transitions between clusters in the training data sequence with *p* = 0.2 (Methods). A neuron learning with LPL still consistently becomes selective to cluster identity even with noisy transitions. **(b)** Cluster selectivity as a function of the probability of noisy cross-cluster transitions in the data sequence with *σ_y_* = 1. LPL drives selectivity to cluster identity only below *p* = 0.5, i.e, only as long as cluster identity remains the slow feature.

**Extended Data Fig. 11:**
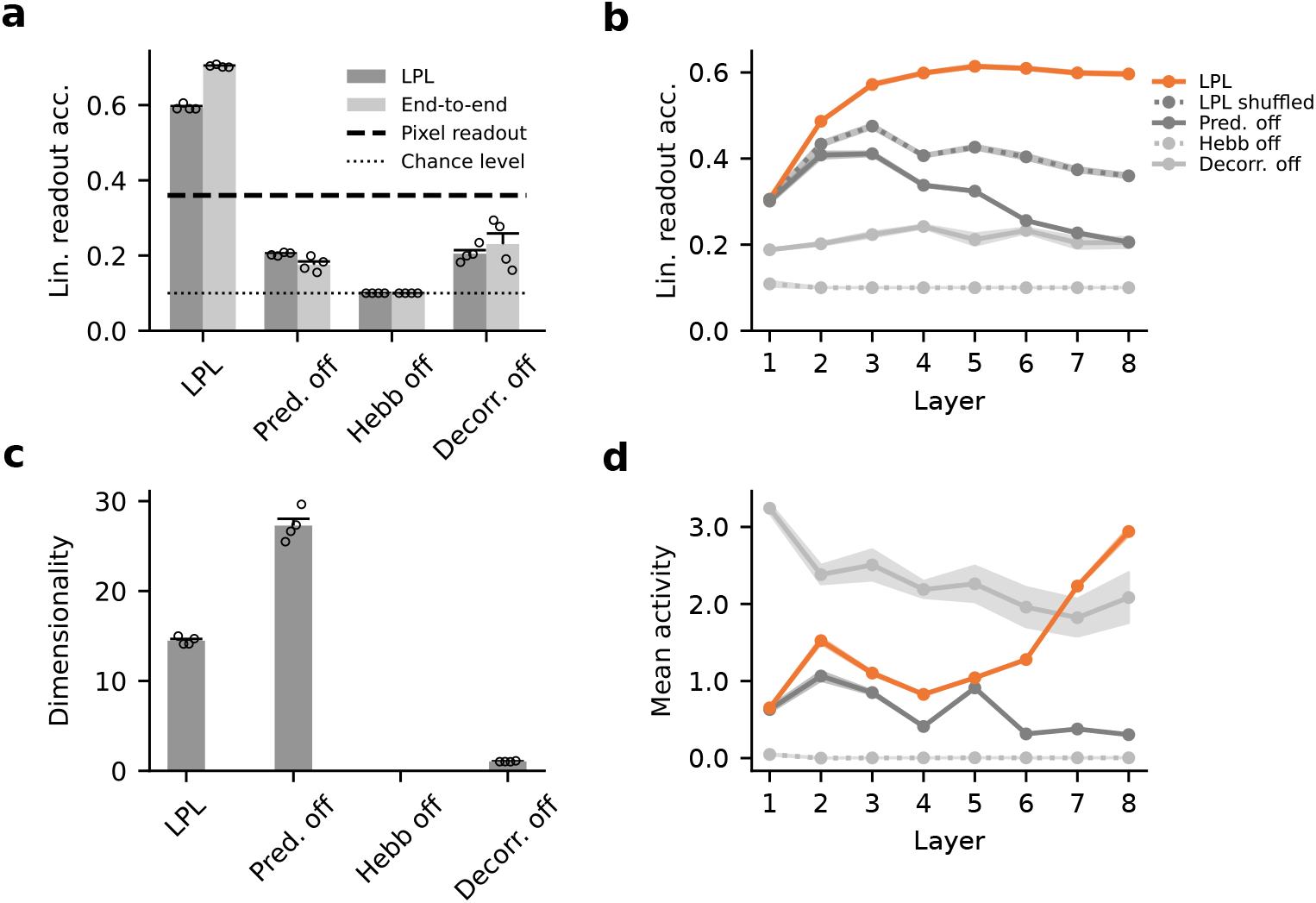
Same as Figure 3 but for the CIFAR-10 dataset. **(a)** Linear readout accuracy of object categories decoded from representations at the network output after training it on natural image data for different learning rules in layer-local (dark) as well as the end-to-end configuration (light). **(b)** Linear readout accuracy of the internal representations at different layers of the DNN after layer-local training. **(c)** Dimensionality of the internal representations for the different learning rule configurations shown in (b). **(d)** Mean neuronal activity at different layers of the DNN after training for the different learning rule variants shown in (b).

**Extended Data Fig. 12:**
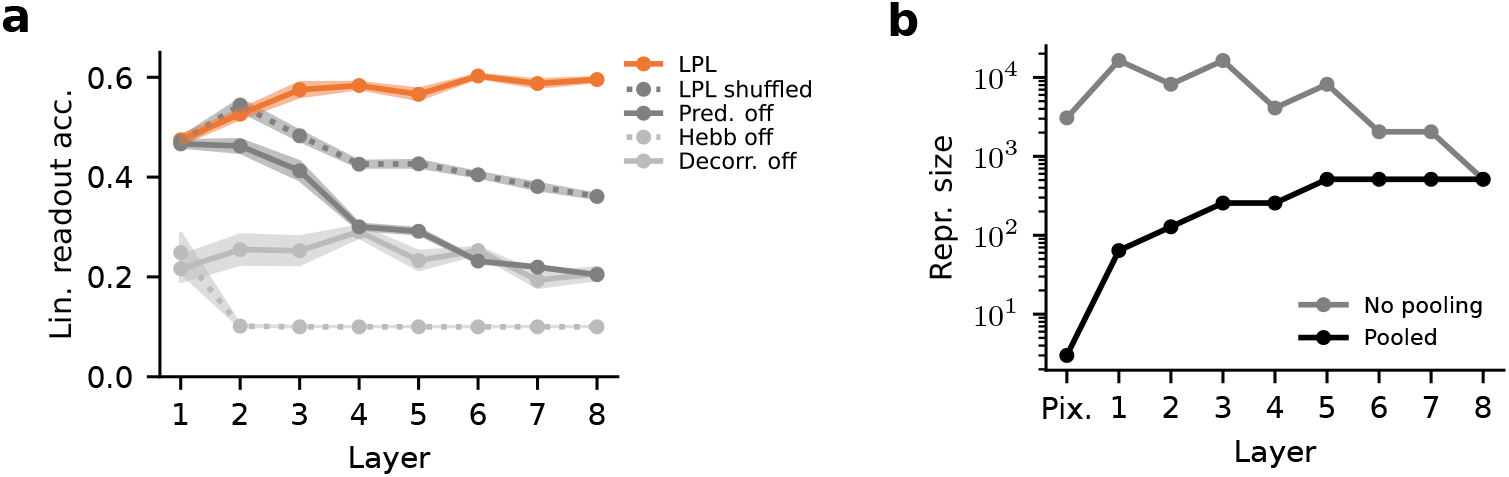
Evaluating readout accuracy without pooling. **(a)** Same as Extended Data Fig. 11b above, but with linear readout accuracies evaluated from the full feature map at each layer instead of after pooling. Results are qualitatively the same as before, but the starting accuracy at early layers is substantially higher. **(b)** Effective representation size that would be the input to the linear classifier at each layer with or without pooling. Without pooling, the number of features at early layers is very large, and may explain the higher early-layer accuracies in (b).

## References

[1] James J. DiCarlo, Davide Zoccolan, and Nicole C. Rust. How Does the Brain Solve Visual Object Recognition? Neuron, 73(3):415–434, 2012-02. ISSN 0896-6273. doi: 10.1016/j.neuron.2012.01.010.

[2] Yann LeCun, Yoshua Bengio, and Geoffrey Hinton. Deep learning. Nature, 521(7553): 436–444, 2015-05. ISSN 0028-0836. doi: 10.1038/nature14539.

[3] Daniel L. K. Yamins, Ha Hong, Charles F. Cadieu, Ethan A. Solomon, Darren Seibert, and James J. DiCarlo. Performance-optimized hierarchical models predict neural responses in higher visual cortex. Proceedings of the National Academy of Sciences, 111(23):8619–8624, 2014-10. ISSN 0027-8424, 1091-6490. doi: 10.1073/pnas.1403112111.

[4] Aran Nayebi, Alexander Attinger, Malcolm Campbell, Kiah Hardcastle, Isabel Low, Caitlin S Mallory, Gabriel Mel, Ben Sorscher, Alex H Williams, Surya Ganguli, et al. Explaining heterogeneity in medial entorhinal cortex with task-driven neural networks. Advances in Neural Information Processing Systems, 34:12167–12179, 2021.

[5] Blake A. Richards, Timothy P. Lillicrap, Philippe Beaudoin, Yoshua Bengio, Rafal Bogacz, Amelia Christensen, Claudia Clopath, Rui Ponte Costa, Archy de Berker, Surya Ganguli, Colleen J. Gillon, Danijar Hafner, Adam Kepecs, Nikolaus Kriegeskorte, Peter Latham, Grace W. Lindsay, Kenneth D. Miller, Richard Naud, Christopher C. Pack, Panayiota Poirazi, Pieter Roelfsema, João Sacramento, Andrew Saxe, Benjamin Scellier, Anna C. Schapiro, Walter Senn, Greg Wayne, Daniel Yamins, Friedemann Zenke, Joel Zylberberg, Denis Therien, and Konrad P. Kording. A deep learning framework for neuroscience. Nature Neuroscience, 22(11):1761–1770, 2019. ISSN 15461726. doi: 10.1038/s41593-019-0520-2.

[6] SueYeon Chung and L. F. Abbott. Neural population geometry: An approach for understanding biological and artificial neural networks. Current Opinion in Neurobiology, 70: 137–144, 2021-10. ISSN 0959-4388. doi: 10.1016/j.conb.2021.10.010.

[7] E. L. Bienenstock, L. N. Cooper, and P. W. Munroe. Theory of the development of neuron selectivity: orientation specificity and binocular interaction in visual cortex. J. Neurosci., 2:32–48, 1982. doi: 10.1523/JNEUROSCI.02-01-00032.1982.

[8] Leon N. Cooper and Mark F. Bear. The BCM theory of synapse modification at 30: interaction of theory with experiment. Nature Reviews Neuroscience, 13(11):798–810, 2012-01. ISSN 1471-003X. doi: 10.1038/nrn3353.

[9] Jordan Guerguiev, Timothy P. Lillicrap, and Blake A. Richards. Towards deep learning with segregated dendrites. eLife, 6:e22901, 2017-12. ISSN 2050-084X. doi: 10.7554/eLife.22901.

[10] João Sacramento, Rui Ponte Costa, Yoshua Bengio, and Walter Senn. Dendritic cortical microcircuits approximate the backpropagation algorithm. Advances in Neural Information Processing Systems, 31, 2018.

[11] Timothy P. Lillicrap, Adam Santoro, Luke Marris, Colin J. Akerman, and Geoffrey Hinton. Backpropagation and the brain. Nature Reviews Neuroscience, pages 1–12, 2020-04. ISSN 1471-0048. doi: 10.1038/s41583-020-0277-3. Publisher: Nature Publishing Group.

[12] Alexandre Payeur, Jordan Guerguiev, Friedemann Zenke, Blake A Richards, and Richard Naud. Burst-dependent synaptic plasticity can coordinate learning in hierarchical circuits. Nature Neuroscience, 24(7):1010–1019, 2021. doi: 10.1038/s41593-021-00857-x.

[13] R P Rao and D H Ballard. Predictive coding in the visual cortex: A functional interpretation of some extra-classical receptive-field effects. Nature Neuroscience, 2(1):79–87, 1999-01. doi: 10.1038/4580.

[14] Nuo Li and James J. DiCarlo. Unsupervised Natural Experience Rapidly Alters Invariant Object Representation in Visual Cortex. Science, 321(5895):1502–1507, 2008. doi: 10.1126/science.1160028.

[15] Georg B. Keller and Thomas D. Mrsic-Flogel. Predictive Processing: A Canonical Cortical Computation. Neuron, 100(2):424–435, 2018-10. ISSN 0896-6273. doi: 10.1016/j.neuron.2018.10.003.

[16] Yosef Singer, Yayoi Teramoto, Ben DB Willmore, Jan WH Schnupp, Andrew J King, and Nicol S Harper. Sensory cortex is optimized for prediction of future input. eLife, 7:e31557, June 2018. ISSN 2050-084X. doi: 10.7554/eLife.31557.

[17] Giulio Matteucci and Davide Zoccolan. Unsupervised experience with temporal continuity of the visual environment is causally involved in the development of v1 complex cells. Science Advances, 6(22):eaba3742, 2020. doi: 10.1126/sciadv.aba3742.

[18] Colleen J. Gillon, Jason E. Pina, Jerome Lecoq, Ruweida Ahmed, Yazan Billeh, Shiella Caldejon, Peter Groblewski, Tim M. Henley, India Kato, Eric Lee, Jennifer Luviano, Kyla Mace, Chelsea Nayan, Thuyanh Nguyen, Kat North, Jed Perkins, Sam Seid, Matthew Valley, Ali Williford, Yoshua Bengio, Timothy P. Lillicrap, Blake A. Richards, and Joel Zylberberg. Learning from unexpected events in the neocortical microcircuit. bioRxiv, page 2021.01.15.426915, January 2021. doi: 10.1101/2021.01.15.426915.

[19] Kaiming He, Xinlei Chen, Saining Xie, Yanghao Li, Piotr Dollar, and Ross Girshick. Masked Autoencoders Are Scalable Vision Learners. 2022 IEEE/CVF Conference on Computer Vision and Pattern Recognition (CVPR), pages 15979–15988, June 2022. doi: 10.1109/CVPR52688.2022.01553.

[20] Aaron van den Oord, Yazhe Li, and Oriol Vinyals. Representation Learning with Contrastive Predictive Coding. arXiv:1807.03748 [cs, stat], July 2018. doi: 10.48550/arXiv.1807.03748.

[21] Ting Chen, Simon Kornblith, Mohammad Norouzi, and Geoffrey Hinton. A simple framework for contrastive learning of visual representations. In Hal Daumé III and Aarti Singh, editors, Proceedings of the 37th International Conference on Machine Learning, volume 119 of Proceedings of Machine Learning Research, pages 1597–1607. PMLR, 13–18 Jul 2020.

[22] Laurenz Wiskott and Terrence J. Sejnowski. Slow Feature Analysis: Unsupervised Learning of Invariances. Neural Computation, 14(4):715–770, 2002-04. ISSN 0899-7667. doi: 10.1162/089976602317318938.

[23] Henning Sprekeler, Christian Michaelis, and Laurenz Wiskott. Slowness: An Objective for Spike-Timing–Dependent Plasticity? PLoS Comput Biol, 3(6):e112, 2007-06. doi: 10.1371/journal.pcbi.0030112.

[24] Bernd Illing, Jean Ventura, Guillaume Bellec, and Wulfram Gerstner. Local plasticity rules can learn deep representations using self-supervised contrastive predictions. Advances in Neural Information Processing Systems, 34, 2021.

[25] Lukasz Kusmierz, Takuya Isomura, and Taro Toyoizumi. Learning with three factors: modulating Hebbian plasticity with errors. Current Opinion in Neurobiology, 46:170–177, 2017-09. ISSN 1873-6882. doi: 10.1016/j.conb.2017.08.020.

[26] Adrien Bardes, Jean Ponce, and Yann LeCun. VICReg: Variance-Invariance-Covariance Regularization for Self-Supervised Learning. arXiv:2105.04906 [cs], May 2021. doi: 10.48550/arXiv.2105.04906.

[27] Erkki Oja. Simplified neuron model as a principal component analyzer. Journal of Mathematical Biology, 15(3):267–273, 1982. ISSN 0303-6812. doi: 10.1007/BF00275687.

[28] W. Gerstner and W.M. Kistler. Spiking neuron models. Cambridge University Press New York, 2002.

[29] Per Jesper Sjöström, Gina G Turrigiano, and Sacha B Nelson. Rate, Timing, and Cooperativity Jointly Determine Cortical Synaptic Plasticity. Neuron, 32(6):1149–1164, 2001-12. ISSN 0896-6273. doi: 10.1016/S0896-6273(01)00542-6.

[30] Daniel E. Feldman. The Spike-Timing Dependence of Plasticity. Neuron, 75(4):556–571, 2012-08. ISSN 0896-6273. doi: 10.1016/j.neuron.2012.08.001.

[31] Sindy Löwe, Peter O’Connor, and Bastiaan S. Veeling. Putting an end to end-to-end: Gradient-isolated learning of representations. Advances in Neural Information Processing Systems, 32(NeurIPS), 2019. ISSN 10495258.

[32] Thomas Miconi. Multi-layer hebbian networks with modern deep learning frameworks. arXiv preprint arXiv:2107.01729, 2021. doi: 10.48550/arXiv.2107.01729.

[33] A. Artola, S. Bröcher, and W. Singer. Different voltage-dependent thresholds for inducing long-term depression and long-term potentiation in slices of rat visual cortex. Nature, 347 (6288):69–72, 1990-09. doi: 10.1038/347069a0.

[34] Wickliffe C. Abraham. Metaplasticity: Tuning synapses and networks for plasticity. Nature Reviews Neuroscience, 9(5):387–387, 2008-01. ISSN 1471-003X. doi: 10.1038/nrn2356.

[35] Sukbin Lim, Jillian L. McKee, Luke Woloszyn, Yali Amit, David J. Freedman, David L. Sheinberg, and Nicolas Brunel. Inferring learning rules from distributions of firing rates in cortical neurons. Nature Neuroscience, 18(12):1804–1810, 2015-12. ISSN 1546-1726. doi: 10.1038/nn.4158. Number: 12 Publisher: Nature Publishing Group.

[36] Friedemann Zenke and Wulfram Gerstner. Hebbian plasticity requires compensatory processes on multiple timescales. Philosophical Transactions of the Royal Society B, 372(1715): 20160259, March 2017. ISSN 0962-8436, 1471-2970. doi: 10.1098/rstb.2016.0259.

[37] Peter Földiak. Forming sparse representations by local anti-Hebbian learning. Biological cybernetics, 64(2):165–170, 1990. doi: 10.1007/BF02331346.

[38] Tim P. Vogels, Henning Sprekeler, Friedemann Zenke, Claudia Clopath, and Wulfram Gerstner. Inhibitory Plasticity Balances Excitation and Inhibition in Sensory Pathways and Memory Networks. Science, 334(6062):1569–1573, 2011-12. doi: 10.1126/science.1211095.

[39] Paul D. King, Joel Zylberberg, and Michael R. DeWeese. Inhibitory Interneurons Decorrelate Excitatory Cells to Drive Sparse Code Formation in a Spiking Model of V1. The Journal of Neuroscience, 33(13):5475–5485, 2013-03. ISSN 0270-6474, 1529-2401. doi: 10.1523/JNEUROSCI.4188-12.2013.

[40] David Lipshutz, Charles Windolf, Siavash Golkar, and Dmitri Chklovskii. A biologically plausible neural network for slow feature analysis. Advances in neural information processing systems, 33:14986–14996, 2020.

[41] Li Jing, Pascal Vincent, Yann LeCun, and Yuandong Tian. Understanding dimensional collapse in contrastive self-supervised learning. International Conference on Learning Representations, 2022.

[42] Hyunjik Kim and Andriy Mnih. Disentangling by Factorising. In Andreas Krause Jennifer Dy, editor, Proceedings of the 35th International Conference on Machine Learning, volume 80, pages 2649–2658. PMLR, July 2018.

[43] Yanis Inglebert, Johnatan Aljadeff, Nicolas Brunel, and Dominique Debanne. Synaptic plasticity rules with physiological calcium levels. Proceedings of the National Academy of Sciences, 2020-12. ISSN 0027-8424, 1091-6490. doi: 10.1073/pnas.2013663117. Publisher: National Academy of Sciences Section: Biological Sciences.

[44] Harel Z. Shouval, Mark F. Bear, and Leon N. Cooper. A Unified Model of NMDA Receptor-Dependent Bidirectional Synaptic Plasticity. Proceedings of the National Academy of Sciences, 99(16):10831–10836, June 2002. ISSN 0027-8424, 1091-6490. doi: 10.1073/pnas.152343099.

[45] Jean-Pascal Pfister and Wulfram Gerstner. Triplets of Spikes in a Model of Spike Timing-Dependent Plasticity. The Journal of Neuroscience, 26(38):9673–9682, 2006-09. ISSN 0270-6474, 1529-2401. doi: 10.1523/JNEUROSCI.1425-06.2006.

[46] Claudia Clopath, Lars Büsing, Eleni Vasilaki, and Wulfram Gerstner. Connectivity reflects coding: a model of voltage-based STDP with homeostasis. Nature Neuroscience, 13(3): 344–352, 2010-03. ISSN 1097-6256. doi: 10.1038/nn.2479.

[47] Julijana Gjorgjieva, Claudia Clopath, Juliette Audet, and Jean-Pascal Pfister. A triplet spike-timing–dependent plasticity model generalizes the Bienenstock–Cooper–Munro rule to higher-order spatiotemporal correlations. Proceedings of the National Academy of Sciences, 108(48):19383–19388, 2011-11. doi: 10.1073/pnas.1105933108.

[48] Taro Toyoizumi, Megumi Kaneko, Michael P. Stryker, and Kenneth D. Miller. Modeling the Dynamic Interaction of Hebbian and Homeostatic Plasticity. Neuron, 84(2):497–510, 2014-10. ISSN 0896-6273. doi: 10.1016/j.neuron.2014.09.036.

[49] Michael Graupner and Nicolas Brunel. Calcium-based plasticity model explains sensitivity of synaptic changes to spike pattern, rate, and dendritic location. Proceedings of the National Academy of Sciences, 109(10):3991–3996, 2012-06. ISSN 0027-8424, 1091-6490. doi: 10.1073/pnas.1109359109.

[50] Guillaume Hennequin, Everton J. Agnes, and Tim P. Vogels. Inhibitory Plasticity: Balance, Control, and Codependence. Annual Review of Neuroscience, 40(1):557–579, 2017-07. ISSN 0147-006X. doi: 10.1146/annurev-neuro-072116-031005.

[51] Edmund T. Rolls and Simon M. Stringer. Invariant visual object recognition: A model, with lighting invariance. Journal of Physiology-Paris, 100(1):43–62, July 2006. ISSN 0928-4257. doi: 10.1016/j.jphysparis.2006.09.004.

[52] Yazhe Li, Roman Pogodin, Danica J Sutherland, and Arthur Gretton. Self-supervised learning with kernel dependence maximization. Advances in Neural Information Processing Systems, 34, 2021.

[53] Johannes Mehrer, Courtney J. Spoerer, Emer C. Jones, Nikolaus Kriegeskorte, and Tim C. Kietzmann. An ecologically motivated image dataset for deep learning yields better models of human vision. Proceedings of the National Academy of Sciences, 118(8), 2021-02. ISSN 0027-8424, 1091-6490. doi: 10.1073/pnas.2011417118. Publisher: National Academy of Sciences Section: Biological Sciences.

[54] Jure Zbontar, Li Jing, Ishan Misra, Yann LeCun, and Stephane Deny. Barlow twins: Selfsupervised learning via redundancy reduction. In Marina Meila and Tong Zhang, editors, Proceedings of the 38th International Conference on Machine Learning, volume 139 of Proceedings of Machine Learning Research, pages 12310–12320. PMLR, 18–24 Jul 2021.

[55] Karen Simonyan and Andrew Zisserman. Very deep convolutional networks for large-scale image recognition. arXiv preprint arXiv:1409.1556, 2014.

[56] Ashok Litwin-Kumar, Kameron Decker Harris, Richard Axel, Haim Sompolinsky, and L. F. Abbott. Optimal Degrees of Synaptic Connectivity. Neuron, 93(5):1153–1164.e7, 2017. ISSN 10974199. doi: 10.1016/j.neuron.2017.01.030.

[57] Friedemann Zenke, Everton J. Agnes, and Wulfram Gerstner. Diverse synaptic plasticity mechanisms orchestrated to form and retrieve memories in spiking neural networks. Nature Communications, 6:6922, 2015-04. ISSN 2041-1723. doi: 10.1038/ncomms7922.

[58] Friedemann Zenke and Surya Ganguli. Superspike: Supervised learning in multilayer spiking neural networks. Neural computation, 30(6):1514–1541, 2018. doi: 10.1162/neco_a_01086.

