## Supplementary Information for "The combination of Hebbian and predictive plasticity learns invariant object representations in deep sensory networks"

### Supplementary Tables

| | STL-10 | | CIFAR-10 | |
|---|---|---|---|---|
| | Layer-local (%) | End-to-end (%) | Layer-local (%) | End-to-end (%) |
| LPL | 63.2±0.3 | 72.5±0.1 | 59.4±0.4 | 70.4±0.2 |
| Neg. samples | 77.0±0.2 | 81.0±0.3 | 67.4±0.3 | 76.5±0.1 |
| Supervised | 70.8±0.3 | 77.8.5±0.3 | 81.7±0.2 | 87.1±0.2 |
| Pixel-space decoding | 31.6 | | 35.9 | |

Table S1: Extended version of Table 1 showin linear classification accuracy on the STL-10 and CIFAR-10 datasets for Latent Predictive Learning (LPL) and different baseline models (Methods). Error values correspond to standard error of the mean (SEM) over four simulations with different random seeds.

| | STL-10 | CIFAR-10 |
|---|---|---|
| VGG-11 | 63.2±0.3 | 59.4±0.4 |
| VGG-11 without bias terms | 58.7 | 54.6 |
| VGG-11 with stale mean and variance estimates | 59.3±0.6 | 47.7±0.6 |
| VGG-11 without GAP | 29.2±1.8 | 47.7±0.3 |
| VGG-11 without GAP (down-sampled inputs) | 45.3±0.5 | - |

Table S2: Linear classification accuracy on the STL-10 and CIFAR-10 datasets for layer-locally trained LPL with the base VGG-11 architecture, and several modified versions. VGG-11 without bias terms refers to the standard architecture including GAP but without any bias terms in the convolutional layers. Stale mean and variance estimates denote calculating the representational mean and variance from the previous batch. VGG-11 without GAP refers to computing and optimizing the LPL objective on the unpooled feature maps. Downsampled inputs correspond to the architecture without GAP, but now using STL-10 images subsampled to a lower resolution of $32 \times 32$. Reported error values correspond to SEM over four simulations with different random seeds.

| | Single neuron (2D dataset) | Single neuron (digit dataset) | Network simulations |
|---|---|---|---|
| $\lambda_1$ | 1 | 1 | 1 |
| $\lambda_2$ | - | - | 10 |
| Optimizer | SGD | SGD | Adam[a] |
| Learning rate $\eta$ | $\min(10^{-2}, 10^{-2}/\sigma_y)$ | $10^{-2}$ | $10^{-3}$ [b] |
| Weight decay $\eta_w$ | 0.15 | 0.15 | $1.5 \times 10^{-6}$ |
| Batch size B | 200 | 200 | 1024 |
| Training steps | $\max(10000, 100\sigma_y)$ | 1000 | ∼78000[c] |

[a] With default parameters from Kingma et al. [1].
[b] Reduced to zero during training using a cosine learning rate schedule.
[c] ∼35000 for CIFAR-10.

Table S3: Hyperparameter values for deep neural network (DNN) simulations.

| Parameter | Value | Parameter | Value |
|---|---|---|---|
| $\tau^{\text{mem}}$ | 20 ms | $U^{\text{exc}}$ | 0 mV |
| $\tau^{\text{ampa}}$ | 5 ms | $U^{\text{inh}}$ | -80 mV |
| $\tau^{\text{gaba}}$ | 10 ms | $U^{\text{leak}}$ | -70 mV |
| $\tau^{\text{nmda}}$ | 100 ms | | |
| $\tau^{\text{thr}}$ | 5 ms | | |

Table S4: Table summarizing the neuronal parameters.

| Source | Destination | Connection probability | Initial weight value |
|---|---|---|---|
| input | exc | 0.1 | 0.15 |
| exc | inh | local (see Methods) | 0.4 |
| inh | exc | 0.5 | 0.1 |
| inh | inh | 0.1 | 0.4 |

Table S5: Summary of spiking neural network (SNN) connectivity parameters.

### Supplementary Notes

#### S1 Equivalence of the objective function and learning rule formulations

Here, we show that the objective functions defined in Eqs. (4), (5), and (6) indeed result in the LPL rule (Eq. (7)).

**Predictive component.** We recall that the predictive objective $\mathcal{L}_{\text{pred}}$ is the mean squared difference between neuronal activity in consecutive time steps.

$$\mathcal{L}_{\text{pred}} = \frac{1}{2MB}\sum_{b=1}^{B}\|z^b - SG(z^b(t-\Delta t))\|^2 = \frac{1}{2MB}\sum_{b=1}^{B}\sum_{i=1}^{M}\Big(z_i^b - SG(z_i^b(t-\Delta t))\Big)^2$$

Taking the derivative with respect to the network weights results in the following learning rule

$$\frac{\partial \mathcal{L}_{pred}}{\partial W_{ij}} = \frac{1}{MB}\sum_{b=1}^{B}\Big(z_i^b - z_i^b(t-\Delta t)\Big) f'(a_i^b)x_j^b \tag{1}$$

which does not require backpropagation through time due to the Stopgrad function.

**Hebbian component.** The Hebbian component minimizes the negative logarithm of the variance of neuronal activity:

$$\mathcal{L}_{Hebb} = \frac{1}{2M}\sum_{i=1}^{M} -\log\left(\sigma_i^2\right)$$

where $\bar{z}_i = \text{SG}(\frac{1}{B}\sum_{b=1}^{B} z_i^b)$ and $\sigma_i^2 = \frac{1}{B-1}\sum_{b=1}^{B}\left(z_i^b - \bar{z}_i\right)^2$ are the mean and variance of the activity of the $i$th output neuron over the minibatch. The corresponding learning rule is obtained as the negative gradient of this loss function with respect to the weights $W$. The gradient itself is given by:

$$\frac{\partial \mathcal{L}_{Hebb}}{\partial W_{ij}} = -\frac{1}{M(B-1)\sigma_i^2}\sum_{b=1}^{B}(z_i^b - \bar{z}_i)f'(a_i^b)x_j^b \tag{2}$$

Note that the objective and the resulting gradient is essentially unchanged upto a scaling factor when we use running estimates of the variance with a time constant $\tau$ instead of batch estimates.

$$\begin{aligned}
\mathcal{L}_{Hebb}(t) &= \frac{1}{2M}\sum_{i=1}^{M} -\log\left(\sigma_i^2(t)\right) \\
&= \frac{1}{2M}\sum_{i=1}^{M} -\log\left((1-\tau)(z_i(t)-\bar{z}_i(t))^2 + \tau\text{SG}(\sigma_i^2(t-1)) + \epsilon\right) \\
\frac{\partial \mathcal{L}_{Hebb}}{\partial W_{ij}}(t) &= -\frac{(1-\tau)}{M\sigma_i^2(t)}(z_i^b(t) - \bar{z}_i(t))f'(a_i^b(t))x_j^b(t)
\end{aligned}$$

**Decorrelation component.** Finally, the decorrelation objective is the decorrelation loss function as the sum of the squared off-diagonal terms of the covariance matrix between units.

$$\mathcal{L}_{\text{decorr}} = \frac{1}{2(B-1)(M^2-M)}\sum_{b=1}^{B}\sum_{i=1}^{M}\sum_{k\neq i}(z_i^b - \bar{z}_i)^2(z_k^b - \bar{z}_k)^2$$

which gives the gradient:

$$\frac{\partial \mathcal{L}_{\text{decorr}}}{\partial W_{ij}} = \frac{1}{(B-1)(M^2-M)} \sum_{b=1}^{B} (z_i^b - \bar{z}_i) f'(a_i^b) x_j^b \sum_{k \neq i} (z_k^b - \bar{z}_k)^2 \tag{3}$$

**The full learning rule.** The combined weight updates for a descent along the sum of the three gradients in Eqs. (1), (2), and (3) in a single-layer network finally yields the LPL rule including the decorrelation component:

$$\begin{aligned} \Delta W_{ij} &= -\eta \left( \frac{\partial \mathcal{L}_{\text{pred}}}{\partial W_{ij}} + \lambda_1 \frac{\partial \mathcal{L}_{\text{Hebb}}}{\partial W_{ij}} + \lambda_2 \frac{\partial \mathcal{L}_{\text{decorr}}}{\partial W_{ij}} \right) \qquad (4) \\ &= \frac{\eta}{MB} \sum_{b=1}^{B} \left( -(z_i^b - z_i^b(t-\Delta t)) + \lambda_1 \frac{\alpha}{\sigma_i^2} (z_i^b - \bar{z}_i) - \lambda_2 \beta (z_i^b - \bar{z}_i) \sum_{k \neq i} (z_k^b - \bar{z}_k)^2 \right) f'(a_i^b) x_j^b \end{aligned}$$

where $\alpha = \frac{B}{B-1}$ and $\beta = \frac{B}{(B-1)(M-1)}$ are the appropriate normalizing constants, and $\lambda_1$ and $\lambda_2$ are the loss coefficients. Including weight decay in the weight update finally yields the LPL rule for a network given in Eq. (7).

#### S2 Relating the Hebbian component of LPL to Oja's rule

To see the relation of the Hebbian component of LPL with the classic Oja's rule [2], we consider the case of a single output neuron ($M = 1$), with no nonlinearity ($f'(a) = 1$), along with the assumption that the input is zero-centered ($\bar{x}_j = 0$). Consequently, $\bar{z} = \sum_j W_j \bar{x}_j = 0$ and $\sigma_z^2 = \left\langle (z - \bar{z})^2 \right\rangle = \langle z^2 \rangle$, which yields a very simple Hebbian learning rule for descending the gradient in Eq. (2):

$$\Delta W_j(t) = -\frac{\partial \mathcal{L}_{Hebb}(t)}{\partial W_{ij}} = \frac{z(t) x_j(t)}{\langle z^2 \rangle}$$

This update rule along with a weight decay (with coefficient $\eta_w$) yields a learning rule that, on average, is equivalent to Oja's rule up to a scaling factor, and in fact has exactly the same non-zero fixed point when $\eta_w = 1$, but with different convergence dynamics because of the multiplication by $1/\langle z^2 \rangle$.

$$\begin{aligned}
\Delta W_j(t) &= \frac{z(t) x_j(t)}{\langle z^2 \rangle} - \eta_w W_j \\
\langle \Delta W_j \rangle &= \frac{\langle z x_j \rangle}{\langle z^2 \rangle} - \eta_w W_j \\
&= \frac{1}{\langle z^2 \rangle} \left( \langle z x_j \rangle - \eta_w W_j \langle z^2 \rangle \right) \qquad (5)
\end{aligned}$$

Oja's rule is presented below for reference

$$\begin{aligned}
\Delta W_j^{Oja}(t) &= z(t) x_j(t) - W_j z^2 \\
\left\langle \Delta W_j^{Oja} \right\rangle &= \langle z x_j \rangle - W_j \langle z^2 \rangle \qquad (6)
\end{aligned}$$

### S3 Importance of the variance-dependent modulation of Hebbian learning

To analytically understand the importance of the variance-dependent scaling of the Hebbian term in the learning rule, we first looked at the synthetic two-dimensional learning task from Fig. 2a, and modeled the behaviour of the LPL loss functions under a particular distribution of the representations. Specifically, we considered the case where the two input clusters map to a mixture of two normal distributions in *representation space* with each Gaussian component corresponding to the representations of one input cluster. Each of the two Gaussians are assumed to have a standard deviation of $r$, with their means symmetrically located on either side of zero with a distance of $D = 4r$ between their centers. We used this setting to investigate how the predictive and Hebbian loss terms of LPL behave under different values of the representational variance by co-varying $r$ and $D$.

The predictive loss in this case is proportional to the expected squared difference $\langle (z_1 - z_2)^2 \rangle$ between two independently drawn samples $z_{1/2}$ from the same Gaussian, i.e, $\mathcal{L}_{\text{pred}} = 2r^2$. The overall variance of the representations is $\sigma_z^2 = r^2 + (\frac{D}{2})^2$ (variance of the Gaussian mixture). Under this representational distribution, we studied the overall loss function obtained by combining the predictive loss with different variance-maximising losses, each a different decreasing function of the variance, i.e, $\mathcal{L}_{\text{var}} = f(\sigma_z^2)$. Specifically, we considered the cases $\mathcal{L}_{\text{var}} = -\sigma_z^2$, $\mathcal{L}_{\text{var}} = -\log \sigma_z^2$, and $\mathcal{L}_{\text{var}} = \text{ReLU}(1 - \sigma_z)$. These loss functions are plotted in

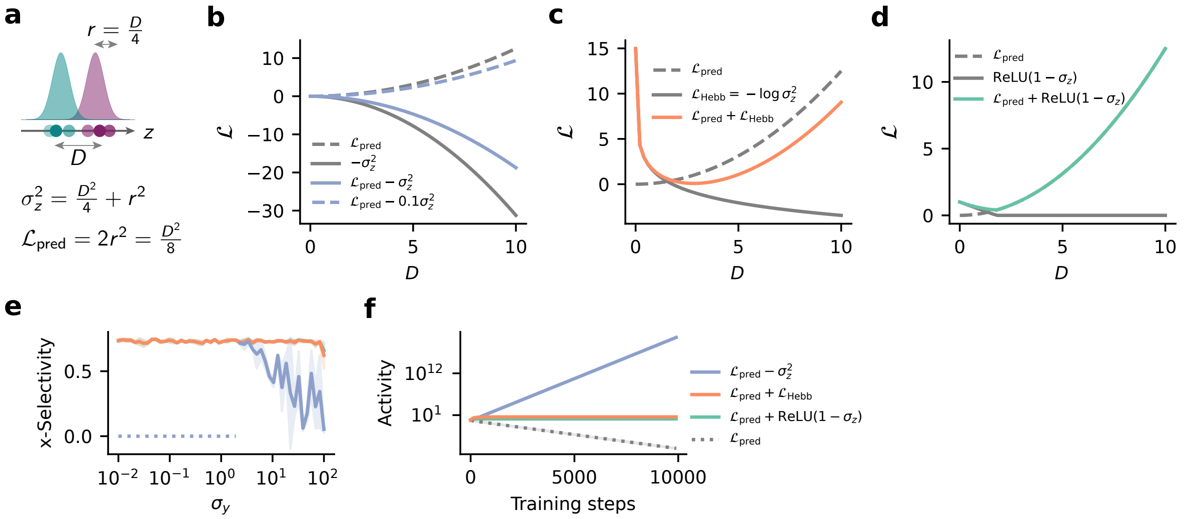


Figure S1: **Variance-dependent modulation of Hebbian learning objective is crucial for stable learning.** **(a)** Example setting with two clusters distance $D$ apart in representation space. The size of each cluster $r = \frac{D}{4}$ is assumed to scale with $D$. In this case, representational variance is $\frac{D^2}{4} + r^2$. **(b)** Total loss combining the predictive loss with a naive variance maximising loss ($-\sigma_z^2$) as a function of cluster separation $D$. Depending on the coefficient of the variance loss, the global minimum of the loss is either at $D = \infty$ (solid blue) or at $D = 0$ (dashed blue), implying runaway long-term potentiation (LTP) or collapse respectively. **(c)** Same as (b) but using $\mathcal{L}_{\text{Hebb}} = -\log \sigma_z^2$, the Hebbian objective optimized by LPL instead of the naive variance objective. Here, the predictive loss starts dominating at higher values of $D$ inducing a global minimum at $D \approx 3$. **(d)** Same as (b) but the variance loss is now $\text{ReLU}(1 - \sigma_z)$, the objective that was optimized in the VICReg model [3]. Here as well, the predictive loss dominates at higher values of $D$, inducing a global minimum at $D \approx 2$. **(e)** Cluster selectivity learned by LPL on the two-dimensional synthetic sequence from Fig. 2a with the standard predictive loss combined with the different variance losses from (a), (b) and (c). Dotted sections indicate simulations where learning diverged. **(f)** Mean output activity over training time for LPL on the two-dimensional synthetic sequence with $\sigma_y = 1$ for the different cases from (a), (b) and (c).

Fig. S1b-d respectively along with $\mathcal{L}_{\text{pred}}$, and the resulting full LPL objective in each case. With the naive variance maximization objective $\mathcal{L}_{\text{var}} = -\sigma_z^2$, the full objective is dominated by the variance term at large values of $D$ (Fig. S1b), and therefore inherently drives unstable learning. It is not possible to remedy this situation by simply using a smaller weight for the variance loss, for instance, by weighting $\mathcal{L}_{\text{var}}$ with a small weight of 0.1. This is because downweighting the variance objective simply moves the loss minimum at $D = \infty$ to $D = 0$, the exact situation of collapse the variance objective is meant to prevent (Fig. S1b). In contrast, using $\mathcal{L}_{\text{var}} = -\log \sigma_z^2$ (Fig. S1c), or $\mathcal{L}_{\text{var}} = \text{ReLU}(1 - \sigma_z)$ (Fig. S1d) along with $\mathcal{L}_{\text{pred}}$ constitute loss landscapes with minima at finite non-zero values of $D$. This is because these variance objectives only dominate at low values of $D$, but have diminishing influence with growing $D$ allowing the predictive term to dictate learning, and preventing runaway activity.

We validated these scaling arguments with learning simulations using each of the three proposed learning objectives on the synthetic two-dimensional sequence learning task from Fig. 2a. We found that using the naive variance maximization objective $\mathcal{L}_{\text{var}} = -\sigma_z^2$ results in poor learning of cluster selectivity, whereas $\mathcal{L}_{\text{var}} = -\log \sigma_z^2$ and $\mathcal{L}_{\text{var}} = \text{ReLU}(1 - \sigma_z)$ prove effective (Fig. S1e). Furthermore, the naive variance objective indeed suffers from runaway instability (Fig. S1f).

### S4 Predictive feature selected by LPL strictly depends on temporal contiguity properties of the input sequence

The slow or "predictive" feature picked up by a single neuron learning with LPL purely depends on the temporal order of stimuli that it is exposed to. One would expect, then, that it is possible to manipulate the learned feature by altering the temporal sequence of stimuli.

To illustrate that this is indeed the case, we designed a predictive learning task similar to that in Fig. 2a using a subset of images from the MNIST handwritten digit dataset [4]. Specifically, we sampled 2000 images from this dataset corresponding to the digits "five" and "six", equally distributed between the two classes. We generated sample inputs by embedding these $28 \times 28$ grayscale images in a $56 \times 56$ blank canvas at either the top-left or bottom-right location. Because we sought to demonstrate that changing the temporal transition structure qualitatively changes neuronal selectivity, we considered two types of sequences with distinct temporal contiguity properties (Fig. S2). In the Digit Sequence we preserved digit identity (either five or six) across subsequent input images, while changing their position on the canvas, whereas in the Location Sequence, we presented different digits at the same location in successive inputs. Therefore, the predictive feature is location in the Location Sequence and digit identity in the Digit Sequence. Furthermore, digit identity and digit location were approximately aligned with the first two principal components of the data which account for 30 % and 5 % of the explained variance respectively (Fig. S2).

We again exposed a single rate neuron model to these two sequence types, while allowing the plastic input connections to evolve according to the LPL rule. After convergence, we measured neuronal selectivity to digit identity and location. We measured selectivity to location and digit identity with the same measure defined in Eq. (9), only changing what inputs fall into clusters 1 and 2 in each case. Concretely, we measured selectivity to digit identity by setting $\langle z_1 \rangle$ to be the mean response to the digit five (at any location) and $\langle z_2 \rangle$ the mean response to the digit six. Finally, we set $\langle z_1 \rangle$ and $\langle z_2 \rangle$ to the mean responses to digits at the two locations regardless of digit identity in order to measure location selectivity.

At initialization with random weights, the neuron was partially selective to both location and digit identity (Fig. S2f). However, subsequent training with LPL rendered the neuron purely selective to either location or digit identity depending on which sequence it was exposed to during training. Yet, when the predictive term was turned off, the specific sequence did not matter and the neuron always became selective to location, which coincides with the direction of highest variance in the data (PC1; Supplementary Fig. S2). Finally, we confirmed that Oja's rule showed the same behavior (Fig. S2f). Thus, a neuron learning with LPL finds temporally contiguous features in high-dimensional sequential data rather than the direction of largest variance, and the the temporally contiguous feature that is learned is strictly determined by the temporal sequence of the stimuli the neuron is exposed to.

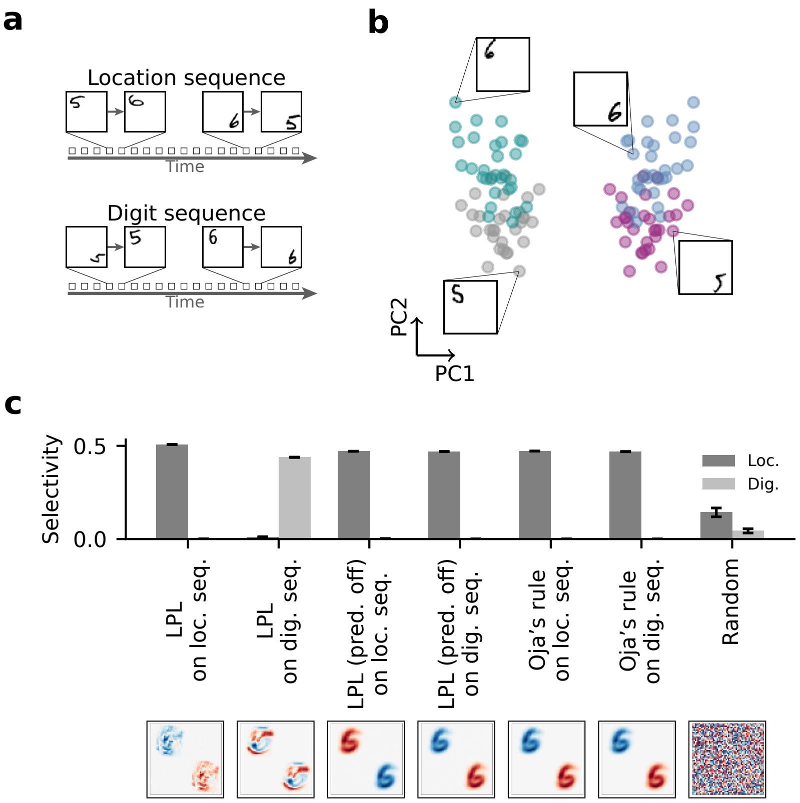


Figure S2: **Temporal contiguities determine features learned under LPL.** **(a)** Schematic of the two kinds of temporal sequences (Methods) in which subsequent inputs were either different digits shown at the same location ("Location Sequence") or the same digit presented at a different location ("Digit Sequence"). **(b)** Scatter plot of the first two principal components of the synthetic digit dataset consisting of randomly sampled handwritten digits in one of the two locations. The principal components closely correspond to location (PC1) and digit identity (PC2). The four digit-position categories are indicated by color. Insets show representative examples from each category. **(c)** Emergent feature selectivity of a single neuron exposed to the two sequence modalities while learning under different rules (top), and the resulting input weights learned in each case (bottom). Under LPL, the neuron's selectivity mirrors the temporally preserved (predictive) property in each sequence, i.e., location in the Location Sequence and digit identity in the Digit Sequence. However, the specific sequence does not matter for Oja's rule or for LPL without the predictive term. In these cases, the neuron always becomes selective to location, the direction of maximum variance. Error bars indicate SEM over ten random seeds.

### S5 Details of the deep neural network architecture

For all DNN simulations, we used the convolutional layers of the VGG-11 architecture consisting of eight blocks each containing $3 \times 3$ convolutions, the ReLU activation function followed by a $2 \times 2$ max-pool operation in some blocks (detailed architecture description provided below).

```
VGG11Encoder(
  (blocks): ModuleList(
    (0): ConvBlock(
      (module): Sequential(
        (0): Conv2d(3, 64, kernel_size=(3, 3), stride=(1, 1), padding=(1, 1))
        (1): ReLU(inplace=True)
        (2): MaxPool2d(kernel_size=2, stride=2, padding=0, dilation=1, ceil_mode=False)
      )
    )
    (1): ConvBlock(
      (module): Sequential(
        (0): Conv2d(64, 128, kernel_size=(3, 3), stride=(1, 1), padding=(1, 1))
        (1): ReLU(inplace=True)
        (2): MaxPool2d(kernel_size=2, stride=2, padding=0, dilation=1, ceil_mode=False)
      )
    )
    (2): ConvBlock(
      (module): Sequential(
        (0): Conv2d(128, 256, kernel_size=(3, 3), stride=(1, 1), padding=(1, 1))
        (1): ReLU(inplace=True)
        (2): Identity()
      )
    )
    (3): ConvBlock(
      (module): Sequential(
        (0): Conv2d(256, 256, kernel_size=(3, 3), stride=(1, 1), padding=(1, 1))
        (1): ReLU(inplace=True)
        (2): MaxPool2d(kernel_size=2, stride=2, padding=0, dilation=1, ceil_mode=False)
      )
    )
    (4): ConvBlock(
      (module): Sequential(
        (0): Conv2d(256, 512, kernel_size=(3, 3), stride=(1, 1), padding=(1, 1))
        (1): ReLU(inplace=True)
        (2): Identity()
      )
    )
    (5): ConvBlock(
      (module): Sequential(
        (0): Conv2d(512, 512, kernel_size=(3, 3), stride=(1, 1), padding=(1, 1))
        (1): ReLU(inplace=True)
        (2): MaxPool2d(kernel_size=2, stride=2, padding=0, dilation=1, ceil_mode=False)
      )
    )
    (6): ConvBlock(
      (module): Sequential(
        (0): Conv2d(512, 512, kernel_size=(3, 3), stride=(1, 1), padding=(1, 1))
        (1): ReLU(inplace=True)
        (2): Identity()
      )
    )
    (7): ConvBlock(
      (module): Sequential(
        (0): Conv2d(512, 512, kernel_size=(3, 3), stride=(1, 1), padding=(1, 1))
        (1): ReLU(inplace=True)
        (2): MaxPool2d(kernel_size=2, stride=2, padding=0, dilation=1, ceil_mode=False)
      )
```

```
    )
  )
  (pooler): AdaptiveAvgPool2d(output_size=(1, 1))
)
```

Furthermore, for the simulations modeling unsupervised learning in the inferotemporal cortex (IT), we used an adaptive average pooling layer with spatial output dimensions of $13 \times 1$. This ensured that the final pooling layer preserved spatial separation along the canvas itself so that the final feature map consisted of $13 \times 1$ 512-dimensional vectors. We added a fully connected layer on top of these feature maps to finally get a single 512-dimensional feature vector per image.

#### References


[1] Kingma, D. and Ba, J. "Adam: A Method for Stochastic Optimization". In: *arXiv:1412.6980 [cs]* (2014). arXiv: 1412.6980.

[2] Oja, E. "Simplified neuron model as a principal component analyzer". In: *Journal of Mathematical Biology* 15.3 (1982), pp. 267–273. ISSN: 0303-6812. DOI: 10.1007/BF00275687.

[3] Bardes, A., Ponce, J., and LeCun, Y. "VICReg: Variance-Invariance-Covariance Regularization for Self-Supervised Learning". In: *arXiv:2105.04906 [cs]* (May 2021). DOI: 10.48550/arXiv.2105.04906.

[4] LeCun, Y., Bottou, L., Bengio, Y., and Haffner, P. "Gradient-based learning applied to document recognition". In: *Proceedings of the IEEE* 86.11 (1998), pp. 2278–2324.
